# Thermal preconditioning modulates coral physiology and heat tolerance: A multi-species perspective

**DOI:** 10.1101/2024.07.18.604102

**Authors:** Erik F. Ferrara, Anna Roik, Franziska Wöhrmann-Zipf, Maren Ziegler

## Abstract

Global warming threatens reef-building corals by challenging their natural adaptive capacity. Therefore, interventions such as stress hardening by thermal preconditioning could become crucial for their survival. Stress-hardening approaches recognize that organisms living in thermally variable environments are better able to withstand marine heat waves. However, a systematic assessment of preconditioning effects on the baseline physiology and thermal tolerance across coral species is lacking. We assessed the changes of thermal tolerance in six stony coral species (*Galaxea fascicularis, Porites rus, Acropora muricata, Montipora digitata*, and *Stylophora pistillata*) in three thermal preconditioning treatments of stable-high 29 °C and variable-high 29 °C with a daily oscillation of ± 1.5 °C, compared to corals in stable-ambient 26 °C. We quantified changes in photosynthetic efficiency and coral bleaching intensity before and after a short-term heat stress assay and up to 30 days later. Stress-hardening success after preconditioning was observed in nearly all preconditioned corals, but the increases in thermal tolerance were species-specific. The greatest increase was recorded in *G. fascicularis* and *A. muricata*, with stress responses reduced by over 80 %. In contrast, preconditioning regimes had minor effects on stress tolerance of *S. pistillata*, making it least receptive to this intervention. After 30 days, most stress-hardened species demonstrated higher survival and recovery rates than their conspecifics from the stable-ambient regime. Notably, both preconditioning regimes affected baseline physiology, especially in the branching species, as indicated by minor tissue paling and decreased photosynthetic efficiency. We conclude that implementing thermal stress hardening protocols will require careful consideration of the species-specific receptiveness and evaluation of the potential trade-offs that can be inflicted with the post-conditioning shifts in physiological baselines.

## Introduction

Reef-building corals are stenothermal organisms that typically live close to their upper thermal limit (Coles & Jokiel, 1977; Fitt et al., 2001). Therefore, the increasing intensity and frequency of global warming-induced marine heatwaves (Frölicher et al., 2018; Oliver et al., 2018) are the primary drivers of coral bleaching events that threaten the existence of tropical reef ecosystems (Hughes et al., 2018). Coral bleaching, i.e., the breakdown of the obligate symbiosis between the coral host and dinoflagellate microalgae (*Symbiodiniaceae*) (Helgoe et al., 2024), leads to starvation due to the loss of the primary energy source and hence compromises coral health (Coles & Brown, 2003; Fine & Loya, 2002; Wiedenmann et al., 2023).

To continue thriving under changing environmental conditions, corals—like all organisms— rely on adaptive evolutionary mechanisms based on genetic variation and the selection of adaptive alleles (Ellegren & Sheldon, 2008). However, these Darwinian processes of selection act on time scales of generations, so the adaptation of the long-lived corals is currently outpaced by the rapid environmental changes of the last decades (Voolstra & Ziegler, 2020). In addition to such transgenerational adaptation mechanisms, acclimatization responses aid organisms in coping with rapid environmental changes within a lifespan (Hilker et al., 2016). Acclimatization relies on the adjustments of plastic phenotypic traits that can be induced by the changes in gene expression. These allow organisms to reduce stress responses by being better able to cope with extreme environmental conditions without permanent DNA modifications (DeMerlis et al., 2022; Majerova et al., 2021; Roik et al., 2023; Wall et al., 2023; Ziegler et al., 2017). Acclimatization can be initiated through pre-exposure to certain stimuli that “prime” the response to subsequent stronger stimuli or environmental challenges (Ainsworth et al., 2016; Bay & Palumbi, 2015; Middlebrook et al., 2008; T. A. Oliver & Palumbi, 2011; Van Oppen et al., 2015). For instance, corals experiencing seasonal moderate stress events or those living in thermally variable environments such as reef flats and lagoons were reported to be more tolerant to subsequent, more severe thermal extremes compared to conspecifics living in more thermally stable environments (Ainsworth et al., 2016; Bellantuono et al., 2011).

The recognition that corals can develop stress memory through priming stimuli (Barshis et al., 2018; Camp et al., 2017; Drury, 2020; Hackerott et al., 2021; Marhoefer et al., 2021; Marzonie et al., 2023; Schoepf et al., 2015, 2020) is the conceptual foundation for so-called “stress-hardening approaches”, which aim to help accelerate acclimatization of reef-building corals to future ocean warming scenarios (Van Oppen et al., 2015). Implementing stress-hardening concepts in restoration efforts promises to improve the efficiency of such interventions.

The nature of the stimulus is crucial for the priming outcome, as its effects are likely dose-dependent, and excessively strong stimuli may even be detrimental to the organism (Brown et al., 2023; Hackerott et al., 2021; Martell, 2023). These intricacies are reflected by several recent studies which documented only minor or negligible positive effects (Barshis et al., 2018; Henley et al., 2022) or, in some cases, detrimental outcomes (Klepac & Barshis, 2020; Putnam & Edmunds, 2011; Schoepf et al., 2019; Thomas et al., 2018). Further, fine-scale variations in the characteristics of a thermal priming stimulus, such as the diel temperature amplitude or the absolute temperature increase, can significantly alter the outcomes of preconditioning regimes within and across coral species (Calabrese et al., 2007; Calabrese & Mattson, 2017; Carelli & Iavicoli, 2002). For instance, the thermal tolerance of corals and the amplitude of diel temperature fluctuations followed a parabolic relationship, where the intermediate amplitudes of 2–3 °C maximized stress tolerance (Brown et al., 2023), while in other studies, larger amplitudes above 4 °C led to detrimental effects (Klepac & Barshis, 2022; Martell, 2023; Schoepf et al., 2019). Regarding stress-hardening success, of stimulus of thermal variability has generally outperformed that of a stable temperature (Drury et al., 2022). One explanation suggests that night cooling (a phase in a natural thermal variability regime) could enhance the repair of stress damage accumulated during the daytime when temperatures peak (Bertucci et al., 2015; Kenkel et al., 2015; Rivest et al., 2017). At the same time, the absolute mean temperature of the priming stimulus will also influence the outcome of a preconditioning regime. For example, studies that documented increased thermal tolerance after preconditioning applied a thermal stimulus at least 3 °C above the control (ambient) temperature (Majerová & Drury, 2022; Martell, 2023), whereas a milder increase of 2 °C above ambient was insufficient to increase thermal tolerance in another study (Henley et al. 2022).

The response of corals to thermal preconditioning may also strongly vary between coral species. It is known that species differ in their environmental tolerances, and preliminary evidence from disparate studies suggests they might also exhibit differential receptiveness to environmental priming. For instance, preconditioned *Acropora cervicornis* and *Pocillopora acuta* bleached less than untreated conspecifics when experimentally exposed to heat (DeMerlis et al., 2022; Majerova et al., 2021), while heat stress tolerance of other species, such as *Montipora capitata* did not increase after preconditioning (Henley et al., 2022; Roper et al., 2023). These equivocal results call for a study that systematically compares receptiveness to preconditioning across coral species, as more clarity is urgently needed to refine stress-hardening protocol in support of restoration approaches. Furthermore, the impacts of stress-hardening regimes on the baseline physiological performance of corals remain poorly understood. The physiological changes triggered by thermal preconditioning may, on the one hand, increase the thermal tolerance of corals, but on the other hand, this benefit may come at a cost. For instance, it has been established that after exposure to warmer temperatures, corals were likely to reduce their symbiont cell densities (Cunning et al., 2018; Jurriaans & Hoogenboom, 2020). Such shifts in holobiont functioning can serve acclimatization, but also likely entail physiological trade-offs on other traits, such as growth (Cornwell et al., 2021; Roik et al., 2023; Williamson et al., 2021). Resilience and recovery rates of stress-hardened corals are another knowledge gap, as monitoring following their long-term fate is rare. We know that corals living in habitats that feature natural temperature fluctuations and a higher maximum temperature were not only more tolerant but also recovered more rapidly after heat stress compared to corals living in more stable environments (Oliver & Palumbi, 2011; Schoepf et al., 2015; Thomas et al., 2018). Whether such increased resilience is due to an adaptation process or to the acclimatization response is hard to disentangle in these field studies and without a lab-based experimental setup. To address these several knowledge gaps associated with the stress hardening of corals by environmental priming, this study aimed to systematically assess the receptiveness of six coral species: *Galaxea fascicularis* (Linnaeus, 1767), *Porites rus* (Forskål, 1775), *Acropora muricata* (Linnaeus, 1758), *Montipora digitata* (Dana, 1846), *Pocillopora verrucosa* (Ellis & Solander, 1786), and *Stylophora pistillata* (Esper, 1792) to three thermal preconditioning regimes. For over 18 days, corals were exposed to a “stable-high” (29 °C) and a “variable-high” (29 ± 1.5 °C) thermal regime and compared to corals from a “stable-ambient” (26 °C) control regime and thereafter exposed to an acute heat stress test. The physiology of the corals was monitored throughout the experiment to assess how preconditioning regimes would affect (1) baseline coral physiology after the preconditioning phase, (2) thermal tolerance to an acute heat stress exposure, and (3) the long-term resilience of coral in terms of survival and recovery rates at 30 days after acute heat stress. In addition, we aimed to (4) identify the type of preconditioning regime that is universally most effective across species.

## Materials and Methods

### Experimental overview and coral species

Six stony coral species in four families were selected to investigate the effects of three thermal preconditioning regimes on the baseline physiology, heat stress tolerance, and long-term resilience of corals, comparing to an ambient thermal regime. These species represent the most common reef-building corals found in the Indo-Pacific region. Corals with two distinct growth forms were included, with *Galaxea fascicularis* (Linnaeus, 1767) and *Porites rus* (Forskål, 1775) as massive species, and *Acropora muricata* (Linnaeus, 1758), *Montipora digitata* (Dana, 1846), *Pocillopora verrucos*a (Ellis & Solander, 1786), and *Stylophora pistillata* (Esper, 1792) as branching species. These corals were collected from different reef locations worldwide (Table S7) and cultivated at the *Ocean2100* coral aquarium facility at the Justus Liebig University Giessen, Germany, for 2-6 years before the experiment. In July 2021, 432 coral fragments were produced from four to eight colonies per species (Table S7). Each colony was cut into 12 fragments of ∼3-4 cm in length and maintained in 265 L tanks at 26 °C for at least ten weeks before the start of the experiment. Each tank was connected to a recirculating artificial seawater system with water exchange of 0.7 L/min, water flow of 3-6 cm/s, and a 10:14 light:dark photoperiod with a light intensity of 250 ± 30 μmol photons m^-2^ s^-1^ (measured by Apogee Lightmeter, Model MQ-510). The water temperature of each tank was feedback-controlled (GHL Temp Sensor digital, ProfiLux 3 and 4, GHL Advanced Technology GmbH, Germany, and Schego Heater 300 and 600 W, Schemel & Goetz GmbH, Germany) and recorded every 10 minutes (HOBO MX Pendant Temp, MX2201, Onset, USA). Corals were fed with copepods three days per week (Calanoide Copepoden, Zooschatz, Germany), except during acute heat assays when no food was provided. Due to logistic reasons, the experiment was conducted in three consecutive runs using two coral species at a time: (1) *A. muricata* and *M. digitata*, (2) *P. verrucosa* and *S. pistillata*, (3) *G. fascicularis* and *P. rus*. The experiments were conducted between November 2021 and April 2022. Each experiment consisted of three phases: the preconditioning phase, the acute heat stress test phase, and the recovery phase (Fig. 1).

**Figure 1.**
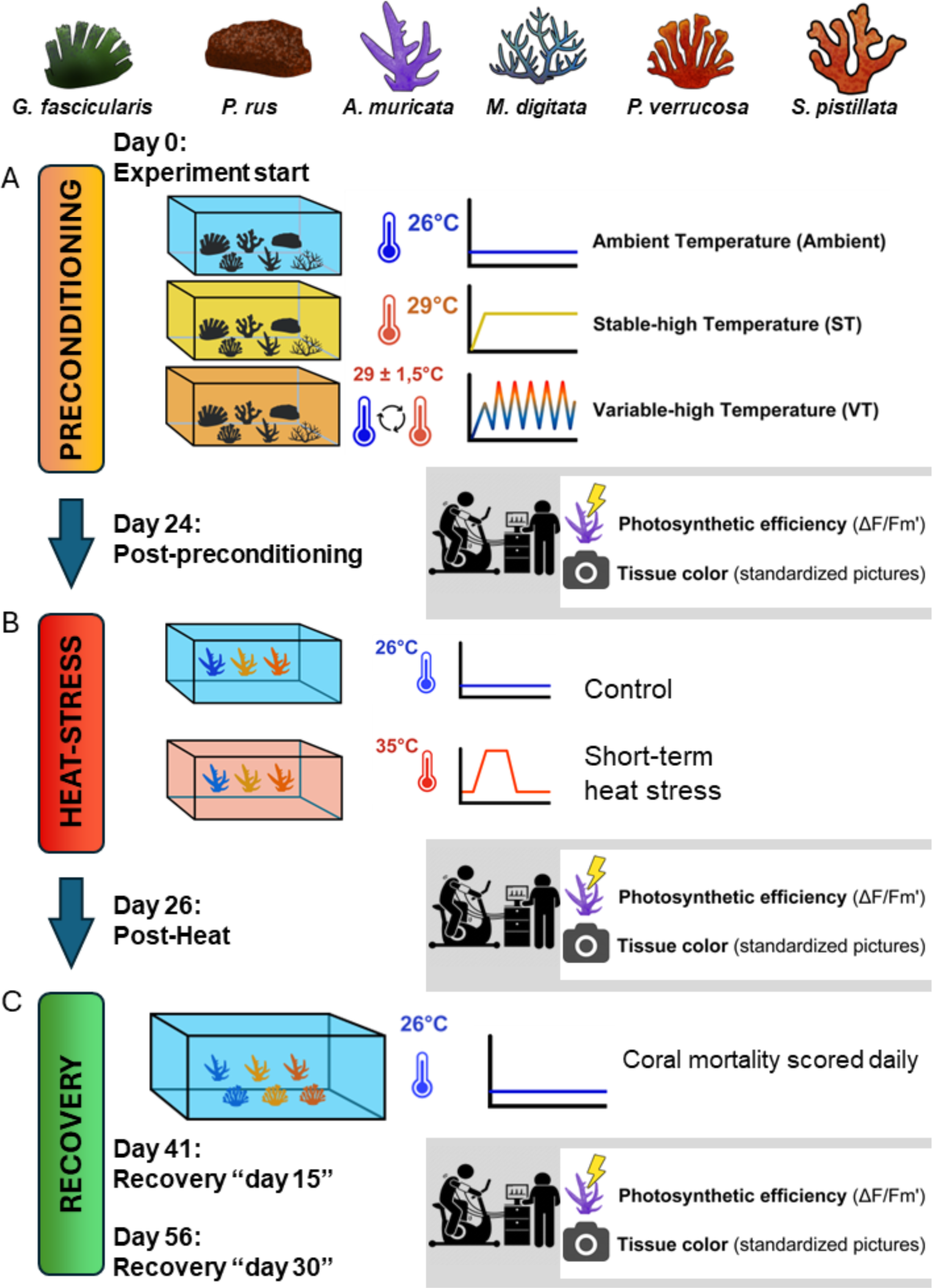
Study design and thermal stress-hardening regimes. Six stony coral species were exposed to three thermal preconditioning regimes (A), followed by short-term heat stress assays (B) to assess the thermal tolerance of corals after preconditioning. Photosynthetic efficiency (effective quantum yield of photosystem II) and tissue color intensity (obtained from standardized pictures) were measured as coral response variables after the preconditioning phase (“post-preconditioning” time point) and after the acute heat stress exposure (“post-heat” time point). Corals were placed in a common tank (C) following the heat stress assay and survivorship was monitored for 30 days. Response variables were measured on days 15 and 30 after heat stress exposure.

### Preconditioning phase

During the preconditioning phase, corals were exposed to three thermal regimes for 24 days, including the stable-high temperature (ST), the variable-high temperature (VT), and a control regime of stable-ambient temperature (Ambient) (Fig. 1A). Twelve fragments per colony were evenly distributed among preconditioning treatment tanks (four fragments per preconditioning regime).

The stable-ambient regime was held at 26 ± 0.5 °C, corresponding to the ambient temperature of the facility. In the stable-high temperature regime, the temperature was increased from 26 °C to 29 °C at 1 °C day^-1^ and kept constant for 18 days (Figure 1 A). The variable-high temperature regime underwent the same temperature increase to 29 °C, after which temperatures fluctuated in a diel cycle with an amplitude of 3 °C around 29 °C for 18 days (Figure 1A). Subsequently, the temperatures in both ST and VT treatments were decreased back to the ambient conditions of 26°C within a day and maintained constant for two more days (lag phase), resulting in a total preconditioning phase of 24 days.

### Heat stress phase

Heat stress assays were conducted after the preconditioning phase (Fig. 1B). The assays were set up using 12 40-L tanks, each equipped with a current pump (easyStream pro ES-28, Aqualight GmbH, Bramsche/Lappenstuhl, Germany) and 65 μm mesh inflow filters to prevent intrusion of particles. The heat stress assays were run at a light intensity of ∼120 μmol photons m^-2^ s^-1^ (white and blue sunaECO LED, AquaRay by Tropical Marine Centre, United Kingdom). For each run, a total of 144 fragments (two species, 12 colonies) were distributed among 12 tanks (six heat and six control), with 12 fragments per tank. Each tank contained three fragments per colony, one from each preconditioning regime of the same species. The control tanks were maintained at 26 °C. In the heat treatment, the temperature was increased from 26 °C to the target temperature of 35 °C in three hours, held at 35 °C for three hours, and then returned to ambient within two hours. Each heat stress assay began at 15:00 h, and following Voolstra et al. (2020), coral responses were assessed 18 hours later at 9:00 am of the following day. For the heat-tolerant species, additional heat stress cycles were performed (Doering et al. 2021), applying the same schedule twice for *P. rus* and three times for *G. fascicularis* and *M. digitata*.

### Recovery phase

Following the short-term heat stress assays, half of the coral fragments were transferred to a shared recovery aquarium and held at 26 °C for 30 days (Fig. 1C), while the other half was frozen for further analyses (Ferrara et al. in preparation) Mortality was scored every day after the heat stress assays until the end of the experiment by recording fragments as dead when no tissue was left on the skeleton. Recovery was monitored by measuring the photosynthetic efficiency and change in tissue color (bleaching) after 15 and 30 days.

### Coral stress response measurements

Physiological response variables were measured for each fragment at the end of the preconditioning and heat stress phases and during and after the recovery period on days 15 and 30. Heat stress phase data presented here always refer to the last heat stress cycle of each species. Our heat-tolerant species required two (*P.rus*) or three (*G. fascicularis* and *M. digitata*) heat stress cycles.

We measured the effective quantum yield of photosystem II (YII) as a proxy of Symbiodiniaceae health using a pulse-amplitude modulation (PAM) fluorometer equipped with a clear plastic tube at the tip of the fiber optic cable to keep a stable distance to the coral surface at a 45 ° angle (PAM-2500 Portable Chlorophyll Fluorometer, Heinz Walz GmbH, Germany).

Tissue color intensity was assessed as a proxy for coral bleaching, where lighter colors indicated a stronger stress response or bleaching severity. Tissue color intensity was obtained from standardized pictures of each coral fragment documented with a digital SLR camera (Nikon D7000) in an evenly illuminated macro photo studio (80 x 80 x 80 cm, Life of Photo). Fragments were placed on a black background with the larger side facing the camera, next to a reference color card for white balance (ColorChecker Passport Photo 2, Calibrite, US). First, the background was removed from each image in Adobe Photoshop 2020. Cropped images were then analyzed with a Python script (Reichert et al., 2021), extracting the gray channel value for each pixel to create a color intensity histogram. The mean value was used to estimate the tissue color intensity. Bleaching severity was assessed from tissue color intensity on a scale of 0 to 255, in which 0 corresponds to white and 255 corresponds to black. While 0 corresponds to pure white, the lowest value reached in completely bleached fragments was 30. Photosynthetic efficiency and tissue color intensity of dead coral fragments during the recovery phase were manually set to 0 and 30, respectively.

### Statistical analyses

All analyses were performed in the *R* statistical environment (version 4.2.3; R Core Team, 2022) with the package *ggplot2* for visualization (Wickham, 2016), *dabestR* v0.3.0 for effect size calculations (Ho et al., 2019), *lme4* v 1.1 - 35.2 for linear mixed effect models (Bates et al., 2015), and *car* for ANOVAs (Fox & Weisberg, 2019). We conducted all analyses separately for each physiological parameter and coral species. Shapiro-Wilk tests were used to test for the normality of the data.

First, the effects of the thermal preconditioning regimes on the baseline physiology of the corals were determined. For this, tissue color intensity and effective photosynthetic yield measured at the end of the preconditioning phase were analyzed separately per species. Differences in physiology between the ST and VT preconditioning regimes compared to the Ambient were calculated as pairwise Hedges’ g effect sizes using raw data. Then, differences were tested with linear mixed effect models applying thermal regimes (ST, 29 °C vs. VT, 29 ± 1.5 °C vs. Ambient, 26 °C) as fixed factor and coral colony genotype as random factor. Normality and homoscedasticity for each model were checked, and where assumptions were violated, data were transformed to improve model performance. ANOVAs were used to compute *F* statistics of the linear mixed effect models.

Next, the effects of the short-term heat stress exposure on the physiology of corals from the three thermal regimes were assessed. Physiological values measured after preconditioning were used as a reference and compared against those measured after heat stress within each preconditioning group with non-parametric Wilcoxon tests. A significant difference indicated the decline of physiological performance due to heat stress exposure. To test whether the preconditioning regime affected the intensity of the heat stress response, we calculated Hedges’ g effect sizes within each preconditioning regime on raw data. Then for each physiological variable for each coral fragment, a heat stress response metric was generated by calculating Δ-values (“post-heat” minus “post-preconditioning” measurements) indicative of the magnitude of the physiological decline due to heat stress. Statistical models compared the differences of Δ-values as proxy for the stress response between the three preconditioning groups using linear mixed effect model with coral colony genotype and tank number as random factors. ANOVAs were used to compute *F* statistics of the linear mixed effect models.

Last, we evaluated whether the effect of heat stress on the physiology was still detectable in the three thermal regimes 30 days after heat exposure. For this, we calculated the effect size as Hedge’s g between control samples not exposed to heat stress and the heat-exposed samples within each preconditioning regime, followed by two-sided permutation t-tests. Survival was scored throughout (1= alive, 0 = dead) and analyzed with the *R* package *survival* (Therneau et al., 2023) and visualized using Kaplan-Meier plots. A heat map of all effect sizes of stress responses, recovery, and survival rates for each coral species was created to summarize and integrate all results into one figure using *ggplot2*.

## Results

### Effects of thermal preconditioning on baseline physiology

Overall, photosynthetic efficiency and tissue color intensity declined in response to both high-temperature preconditioning regimes (Fig. 2). Five of the six tested coral species preconditioned under stable-high temperature (ST, 29 °C) and variable-high temperature (VT, 29 ± 1.5 °C) had significantly lower photosynthetic efficiency and/or tissue color intensity compared to corals in the stable-ambient temperature regime (Ambient, 26°C). Consequently, we classified the six coral species into three main groups based on the magnitude of physiological shifts observed after thermal preconditioning: 1) “robust” for species with stable baseline physiology; 2) “moderately sensitive” for species exhibiting a moderate shift; and 3) “sensitive” for species with large declines in baseline physiology.

**Fig. 2.**
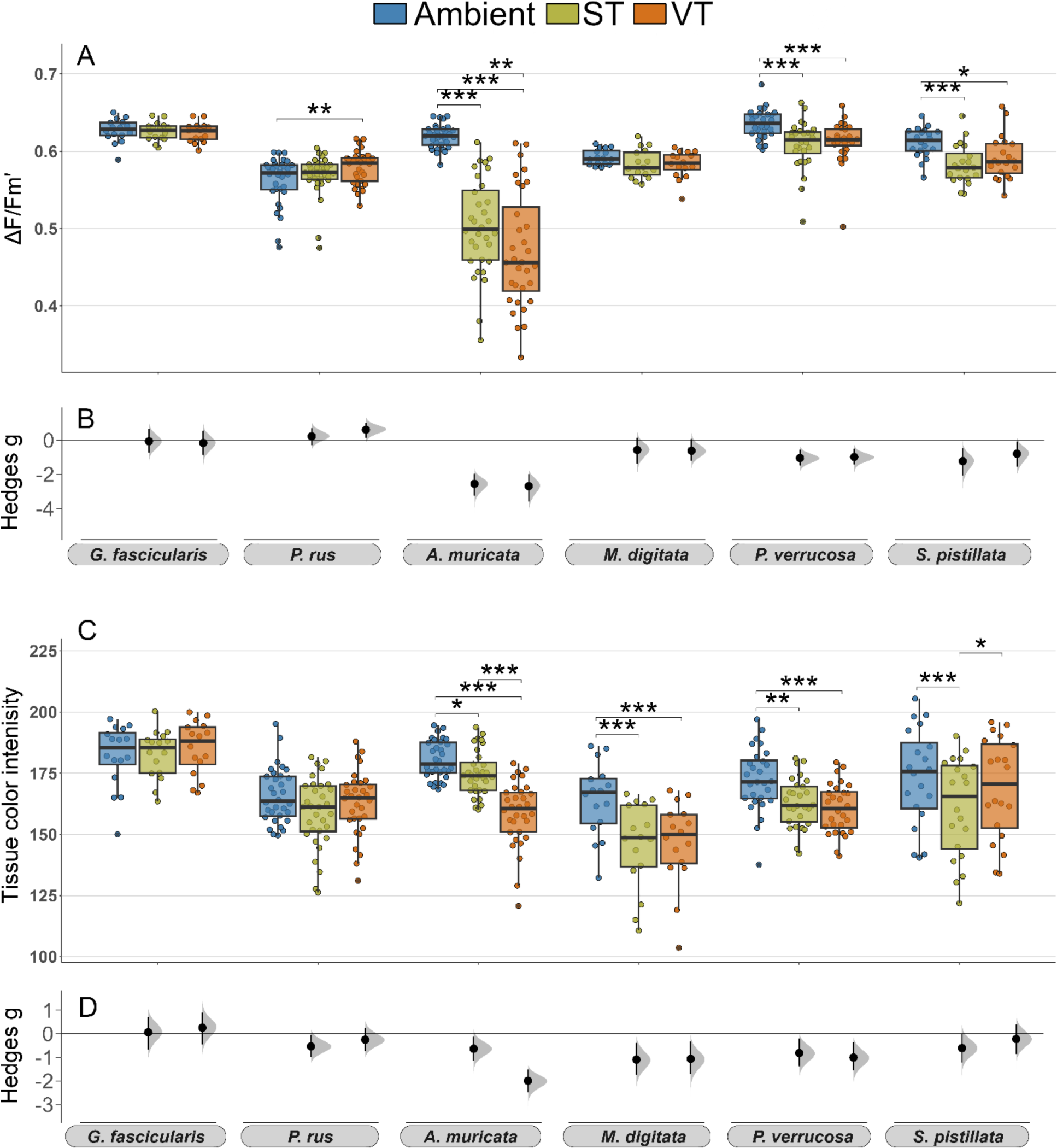
Immediate changes in coral baseline physiology following thermal preconditioning. The changes in effective quantum yield (ΔF/Fm’) (A) and tissue color intensity (C) in response to preconditioning thermal regimes are shown as boxplots, pairwise differences between the stable-high (ST) and variable-high (VT) treatments compared to the stable-ambient (Ambient) treatment are shown as Hedges’ g effect sizes including the 95% CIs (B, D). Data in A and C are displayed as boxplots with raw data points; lines indicate medians, boxes indicate the first and third quartile, and whiskers indicate ± 1.5 IQR. Connecting lines between boxes indicate significant differences between preconditioning regimes (*p* < 0.001***, < 0.01**, < 0.05* from linear mixed effect models).

We classified *Galaxea fascicularis* and *Porites rus* as “robust” species as their photosynthetic efficiency and tissue color intensity remained stable in response to the thermal preconditioning treatments (Fig. 2, Table S1). Specifically, the treatments had no significant effect on the baseline physiology in *G. fascicularis* (*p* > 0.05) and only minor effects in *P. rus,* where photosynthetic efficiency was slightly increased in the VT treatment compared to the Ambient treatment (g = 0.61, *p <* 0.01; Fig. 2A).

We classified *Montipora digitata, Pocillopora verrucosa,* and *Stylophora pistillata* as “moderately sensitive” based on a small but significant decline in their physiological performance in response to thermal preconditioning (Fig. 2). In particular, photosynthetic efficiency significantly decreased by ∼5 % in *P. verrucosa* and *S. pistillata* exposed to the ST and VT preconditioning and compared to those from the Ambient treatment. In contrast, it remained stable in all preconditioning groups for *M. digitata* (*p* > 0.05; Fig. 2A, B, Table S1). Tissue color intensity significantly decreased in ST and VT *M. digitata* and *P. verrucosa* fragments by, on average, 11 and 7 %, respectively (*p* < 0.001 and *p* < 0.01, respectively). Tissue color intensity was significantly reduced in *S. pistillata* ST fragments compared to the Ambient (*p* < 0.001) and VT (*p* < 0.05) groups, which showed similar results (Fig. 2C, D, Table S1).

We ranked *Acropora muricata* as a “sensitive” species based on the comparably large effects of thermal preconditioning on its baseline physiology, which were approximately twice as large as that of the “moderately sensitive” coral species. Photosynthetic efficiency decreased by roughly 20 % in ST and VT corals (*p* < 0.001; Fig. 2A), reaching levels of ΔF/Fm’ below 0.5. Moreover, the decrease in photosynthetic efficiency of VT corals was significantly larger than in ST corals (*p* < 0.01). Similarly, the decrease in tissue color intensity was significantly larger in the VT than in the ST group (*p* < 0.001), which both decreased by 3 and 12 % on average compared to the Ambient group, respectively (ST *p* < 0.05, VT *p* < 0.001; Fig. 2, Table S1). Overall, preconditioned *A. muricata* fragments appeared slightly paler, but visually healthy (Fig. S1b).

### Effects of thermal preconditioning on coral thermal tolerance

Photosynthetic efficiency and tissue color intensity significantly decreased in all coral species in response to the acute heat stress (Fig. 3). However, the heat stress response was consistently more severe in Ambient corals than those from the ST and VT treatments and differences in photosynthetic efficiency were more intense than in tissue color (bleaching).

**Fig. 3.**
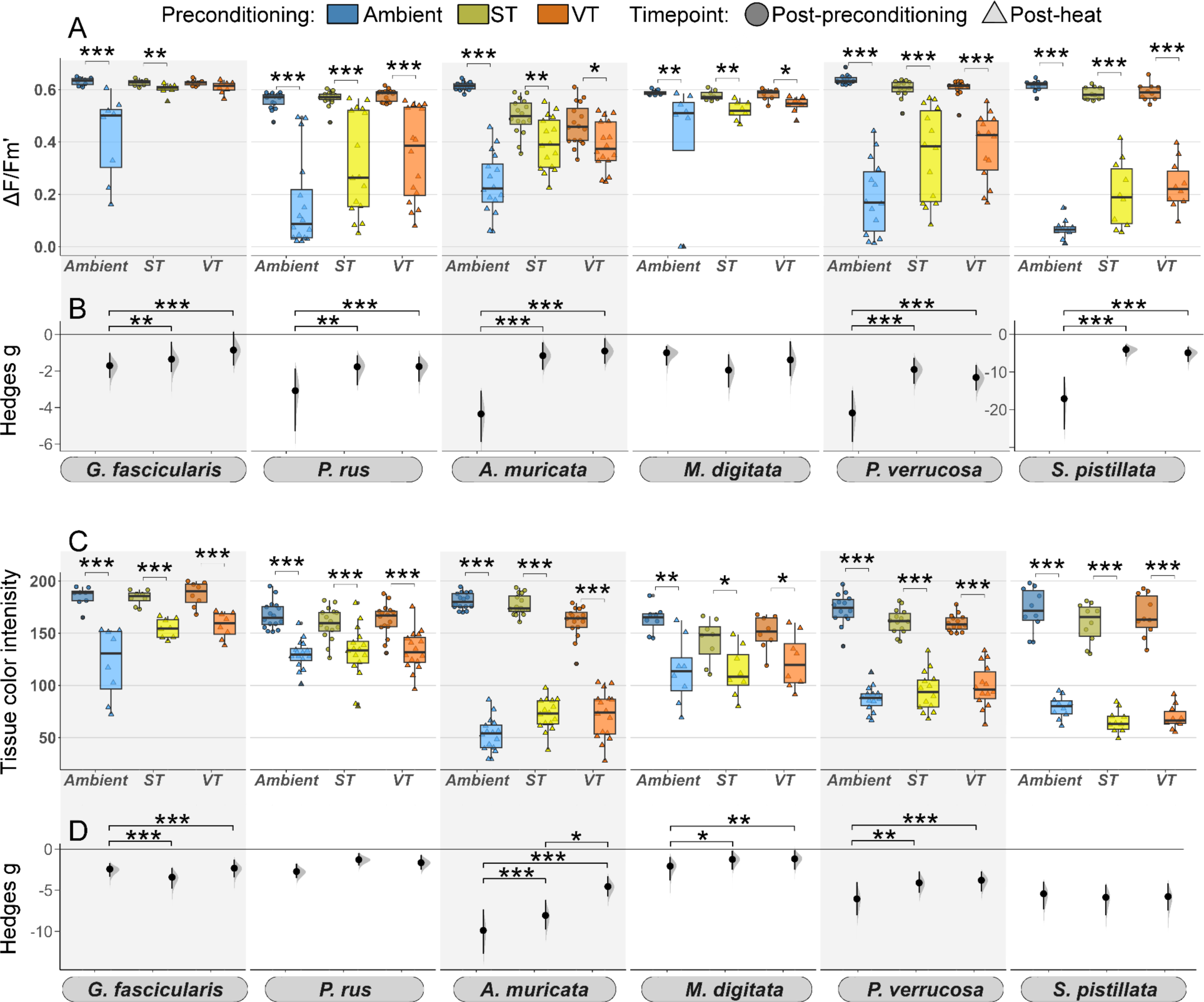
Coral thermal tolerance assessed in heat stress assays following the thermal preconditioning treatments. The decline in effective quantum yield (ΔF/Fm’) (A) and tissue color intensity (C) within each preconditioning group in response to heat stress are shown as boxplots, comparing paired “post-heat” (lighter color) to “post-preconditioning” (darker color) values. Pairwise differences between time points within each preconditioning regime are shown as Cumming estimation plots on Hedges’ g effect sizes, including the 95 % CIs (B, D). Data in A and C are displayed as boxplots with raw data points. Connecting lines between boxes indicate significant differences between time points (*p* < 0.001***, < 0.01**, < 0.05* from Kruskal-Wallis and post hoc Wilcoxon test). Significant differences in heat stress responses between ST and VT compared to the Ambient group are calculated based on the paired delta difference between “post-heat” and “post-preconditioning” time points and indicated by connecting lines (*p* < 0.001***, < 0.01**, < 0.05* from the linear mixed effect model).

We classified the coral species into three groups based on the degree of the stress-hardening effect of preconditioning treatments (ST and VT) on their subsequent response to acute heat stress. We classified coral species as 1) “receptive” to stress-hardening when their heat stress response was significantly smaller in the ST and VT preconditioning groups compared to the Ambient group; 2) as “moderately receptive” when they showed only minor mitigation in stress response and/or responses were inconsistent; 3) and as “not receptive” when the heat stress response was similar across all preconditioning groups.

*Galaxea fascicularis* and *Acropora muricata* were classified as “receptive” to the stress-hardening effect of thermal preconditioning, as their heat stress response was reduced in both response variables (Fig. 3). In *G. fascicularis* and *A. muricata* the declines in photosynthetic efficiency in the ST and VT groups were significantly reduced by 90 and 80 % respectively, compared to the Ambient group (ST *p* < 0.01; VT *p* < 0.001; Fig. 3A, B, Tab. S2, Tab. S4, Tab. S5). Additionally, ST and VT corals bleached significantly less (*p* < 0.001; Fig. 3C, D, Tab. S3, Tab. S4, Tab. S5). In *G. fascicularis*, bleaching of ST and VT corals was half as severe as in the Ambient group (*p* < 0.001; Fig. 3C). While *A. muricata* bleached more severely across all three regimes, the decline in tissue color was significantly smaller in the ST and VT groups compared to the Ambient (*p* < 0.001; Fig. 3C, D). Furthermore, the bleaching severity of VT *A. muricata* fragments was significantly lower than those from the ST group (*p* < 0.5; Fig. 3D). *Porites rus* and *Pocillopora verrucosa* were classified as “moderately receptive” to thermal preconditioning. In these species, the large decrease in photosynthetic efficiency in the Ambient group was mitigated in the ST and ST corals (ST *p* < 0.01; VT *p* < 0.001) while bleaching severity was similar across preconditioning treatments (Fig. 3). The decline in photosynthetic efficiency of ST and VT corals was ∼57% smaller than Ambient *P. rus* and *P. verrucosa* fragments, respectively (Fig. 3A, B, Tab. S2, Tab. S4, Tab. S5). The severity of coral bleaching in response to heat stress was more homogenous across all preconditioning treatments in these “moderately receptive” species than in the “receptive” species (Fig. 3C, D, Tab. S3, Tab. S4, Tab. S5). In *P. rus,* bleaching severity was similar across all preconditioning regimes (*p* > 0.05), while in *P. verrucosa*, ST and VT fragments bleached 30 % less than in the Ambient treatment (ST *p* < 0.01; VT *p* < 0.001; Fig. 3C, D).

Despite the strong difference in inherent heat tolerance between *M. digitata* and *S. pistillata*, both corals were classified as “not receptive” to thermal preconditioning. In both corals, none of the treatments had a relevant effect on the stress responses, however, the specific responses were very different (Fig. 3). In *M. digitata*, the decrease in photosynthetic efficiency after heat stress was minor and the difference between preconditioning treatments was not significant (*p* 0.05; Fig. 3A, B, Tab. S2, Tab. S4, Tab. S5). Additionally, bleaching in *M. digitata* was overall low, with slightly less bleaching in the ST and VT groups compared to the Ambient (*p* 0.01; Fig. 3 C, D). In contrast, photosynthetic efficiency severely decreased across all preconditioning treatments in *S. pistillata,* as illustrated by the largest effect sizes of the heat stress treatment across all species (Fig. 3B) and, although ST and VT preconditioning had a positive effect compared to the Ambient regime (*p* > 0.001), photosynthetic efficiency was extremely low in all groups after heat treatment (i.e., ΔF/Fm’ ∼ 0.2, Fig. 3AB, Tab. S3, Tab. S4, Tab. S5). Moreover, all fragments bleached severely with no difference between the preconditioning treatments (*p* > 0.05; Fig. 3C, D).

### Effects of thermal preconditioning on coral survival and recovery

Coral survival, photosynthetic efficiency, and tissue color were monitored for 30 days after the acute heat stress to evaluate the effect of thermal preconditioning regimes on coral recovery trajectories (Fig. 4). Overall, coral survival was consistently lower in the Ambient group than in the ST and VT groups (Fig. 4A). Moreover, ST and VT corals recovered better. However, survival and recovery rates were species-specific, and recovery dynamics aligned with the inherent thermal tolerance of each coral species.

**Fig. 4.**
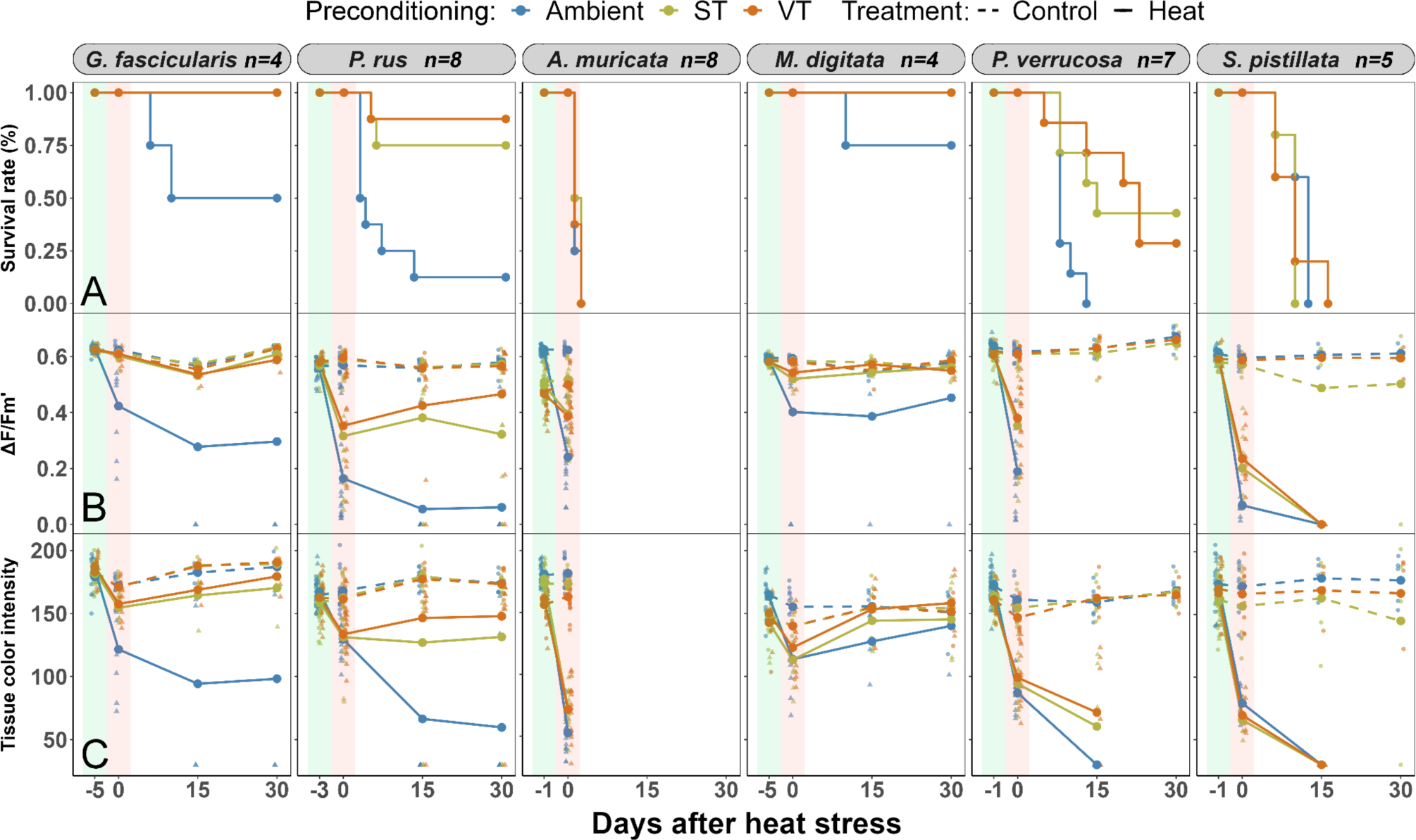
Coral survival rates and recovery following heat stress. Kaplan-Meier plots show changes in coral survival rates under ST, VT, and Ambient preconditioning regimes (A). The trajectories of effective quantum yield (ΔF/Fm’) (B) and tissue color intensity after the heat stress assays (C) for corals from the same preconditioning regimes reflect on the recovery of corals. Lines connect mean values (solid lines for heat treatment, dashed lines for control treatment), raw data points are included. These recovery lines are based on data from a constant sample size within each “population” of each preconditioning regime. The effective quantum yield of dead coral fragments was recorded as 0, while tissue color intensity was scored as 30 (the lowest score observed after the heat stress assay). Physiological parameters of alive *P. verrucosa* fragments were not scored due to uncertainties introduced by algal growth on necrotic tissue, which started dominating the corals between day 0 and 15.

The most heat-tolerant species*, G. fasciculari*s and *M. digitata* exhibited the highest survival rates across all preconditioning treatments. In both species, all ST and VT fragments survived the heat treatment, compared to only 50 and 75 % that survived in the Ambient treatment, respectively (Fig. 4A). Photosynthetic efficiency and tissue color of *G. fascicularis* fragments from the Ambient preconditioning treatment did not entirely recover within the 30 days after heat stress (*p* < 0.05; Fig. 4B, C). In contrast, both metrics recovered to pre-bleaching levels in VT fragments (*p* > 0.05; Fig. 4B, C). In *M. digitata,* photosynthetic efficiency and tissue color of fragments had recovered in all preconditioning groups after 30 days, with slightly slower recovery in corals from the Ambient preconditioning group (p > 0.05; Tab. S3, S6). *P. rus* was the second most heat-stress tolerant species, and it further benefited from the preconditioning, as 75 and 87 % of ST and VT corals survived after heat stress, respectively, compared to 12 % of the Ambient treatment (Fig. 4A). Recovery of photosynthetic efficiency and tissue color was highest in the VT group, where both metrics regained roughly half of the loss immediately after heat stress (*p* > 0.05; Fig. 4B, C). For the ST corals, both metrics remained stable at post-heat stress levels (*p* < 0.05) while they further declined over time in the Ambient group (*p* < 0.001; Tab. S6).

In *P. verrucosa*, all Ambient corals died after heat stress, while 43 and 27 % of ST and VT fragments survived, respectively (Fig. 4a). All surviving fragments had extensive necrotic areas covered by algae, hampering reliable measurements of photosynthetic efficiency and bleaching. In *A. muricata* and *S. pistillata*, which had the lowest inherent heat stress tolerance (Fig. 3), all fragments died before the end of the 30-day monitoring period. This included ST and VT fragments, which initially seemed promising, as heat stress responses were milder (Fig. 4A).

### Comparative evaluation of stress-hardening effects of the thermal preconditioning treatments

In summary, the ST and VT preconditioning treatments had a stress-hardening effect on corals, increasing the thermal tolerance, survival, and recovery rate of most coral species (Fig. 5). Overall, corals preconditioned with the VT treatment exhibited higher thermal tolerance and faster recovery than the ST preconditioned corals (Fig. 5; Tab. S2-S6). This pattern was consistent across most coral species and most metrics. The differences between the effects of the ST and VT treatments were smaller than the difference between these two treatments in comparison to the Ambient treatment, where the strongest declines occurred (Fig. 5). The heat map uses color gradients to summarize and illustrate the species-specific receptiveness to stress-hardening by thermal preconditioning regimes, where *G. fascicularis* was the most receptive species indicated by the overall darker colors in the VT and ST treatments compared to the Ambient, *P. rus* was the species with the highest increase in resilience as indicated by the darker colors in ST and VT during the recovery phase, while *S. pistillata* was the least receptive as indicated by the lighter colors.

**Fig. 5.**
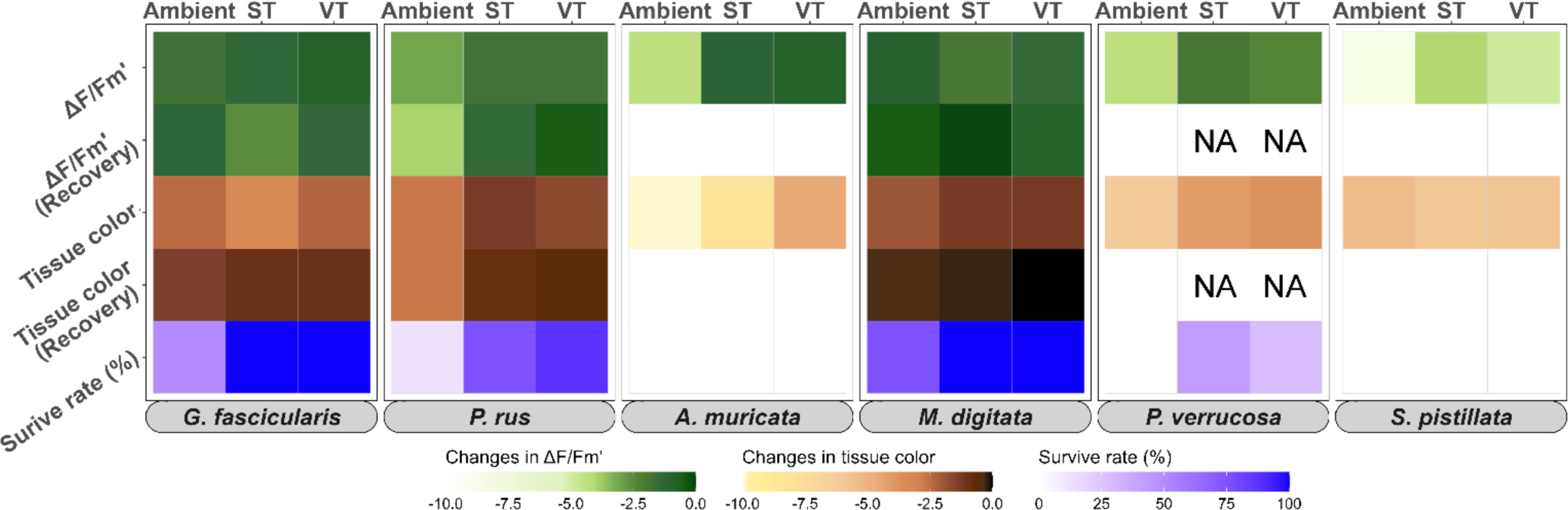
Heat map summarizing stress response physiology, survival, and recovery of six coral species following heat stress. The intensity of change in the measured metrics is coded as intensity of the colors, with lighter shades indicating a more severe response following heat and darker shades indicating stability of the metric. Effect sizes of the heat stress responses (ΔF/Fm’, tissue color) were determined using the Hedges’ g metric as the mean difference between the measurements of the heat and control treatment groups. Values indicate the relative the decrease in photosynthetic efficiency (ΔF/Fm’, green shades) and bleaching score (tissue color, yellow to brown shades) after heat stress and at the end of the recovery period. Kaplan-Meier probabilities represent survival rates (blue to violet shades).

## Discussion

We demonstrated that preconditioning treatments with a stable-high (ST) and a variable-high thermal (VT) regime stress-hardened corals successfully by enhancing their heat tolerance across coral species and, in most cases, by promoting their long-term recovery compared to the untreated corals from the stable-ambient regime (Ambient). The receptiveness to the stress-hardening effect varied across coral species. Additionally, we discovered that the thermal preconditioning regimes triggered a shift in the baseline physiology, resulting in slightly diminished performance in all branching coral species. Our findings highlight the importance of considering coral species traits, such as the receptiveness to stress-hardening and susceptibility to physiological baseline shifts upon the preconditioning (and their potential repercussions) when designing thermal preconditioning protocols for the stress hardening of corals.

### Thermal preconditioning shifts the physiological baselines of corals

Our data revealed that several corals shifted their physiological baseline functioning in response to thermal preconditioning regimes, as we observed a mild decrease in photosynthetic efficiency in the four branching coral species. These corals also appeared paler than those from the Ambient control group, indicating a decrease of symbiont cells, likely reflecting the modulation of symbiont cell densities, which has been long-known to be an acclimatization mechanism of corals along temporal and spatial gradients of temperature and light (Fagoonee et al., 1999; Fitt et al., 2000; Stimson, 1997). Changes in symbiont cell densities are involved in the modulation of heat stress tolerance levels, as for instance, corals with lower symbiont densities are less susceptible to heat stress than those with high symbiont densities (Cornwell et al., 2021; Cunning & Baker, 2013). As symbiont numbers decrease so does the production of hazardous molecules, such as reactive oxygen species during heat stress by these symbionts, linking lower initial symbiont densities to higher resistance and resilience to heat stress (Cornwell et al., 2021; Cunning & Baker, 2013, 2014). This connection could also provide the functional link between the paling of tissues and decline in symbiont photosynthetic efficiency, measured after the ST and VT preconditioning in our study, and the subsequent higher heat stress tolerance.

In contrast to this initial response of the branching corals, physiological parameters of massive-growing corals, *P. rus* and *G. fascicularis,* were less susceptible to both thermal preconditioning treatments. For instance, minor increases rather than decreases in tissue color intensity and photosynthetic efficiency were recorded in *P. rus,* while no changes were recorded in *G. fascicularis*. A naturally low physiological plasticity of these massive corals (Darling et al., 2012) could explain their low susceptibility to the preconditioning treatments. However, massive-growing coral species have previously been shown to modulate symbiont densities in response to changes in temperature and light (Sawall et al., 2022; Ziegler et al., 2015), which demonstrates the capacity to flexibly adjust for physiological needs in certain massive corals. Alternatively, our observation could be explained by their high natural heat tolerance capacity (Jurriaans & Hoogenboom, 2020), allowing them to tolerate the specific ST and VT preconditioning treatments without the need of modifying their physiological functioning. Therefore, it remains to be determined whether these massive species might require and tolerate a more extreme and challenging stimulus than those imposed by our preconditioning treatments. This might result in an even higher increase in thermal tolerance, or it might then become detrimental as observed in other studies (Klepac & Barshis, 2020; Schoepf et al., 2015).

The significant physiological shift in response to the preconditioning treatments in the branching coral species may, on the one hand, provide physiological priming to better cope with heat stress, however, is also likely to entail trade-offs that affect optimal physiological functioning and productivity in the longer term. For instance, the observed reductions in symbiont density and photosynthetic efficiency could be accompanied by reduced skeletal growth rate and energy resource relocation (Cornwell et al., 2021; Roik et al., 2023). A long-term exposure to warmer temperatures can prioritize the relocation of resources into tissue growth while neglecting skeletal growth in reef-building species (Roik et al., 2024) Together with the up-regulation of host metabolism and shifting resource allocation, the energy demands of the holobiont may not be fully covered in the long-term (Gibbin et al., 2018; Rodrigues & Grottoli, 2007; Roik et al., 2023; Wiedenmann et al., 2023). Consequently, the increases in thermal tolerance should be considered a complex trait that will depend on the amount of energy reserves and the energy budgeting strategy of the corals (Grottoli et al., 2006), which will impact the capacity for rapid repair of the tissue damage during heat stress events (Huffmyer et al., 2021). In a worst-case scenario, physiological acclimatization to a short-term extreme condition may ultimately lead to the loss of thermal tolerance in the long-term.

The genetic make-up of symbiont communities can be decisive in shaping photosynthetic performance. Exposure to warmer temperatures may induce shifts in the Symbiodiniaceae community towards assemblages dominated by thermo-tolerant species (e.g., *Durusdinium trenchii*) (Cunning et al., 2018; Silverstein et al., 2015; Williamson et al., 2021). In general, such tolerant symbionts are often characterized by a lower photosynthetic activity at an ambient temperature compared to the less heat-tolerant symbionts (Cantin et al., 2009; Cunning et al., 2018; Quigley et al., 2020; Williamson et al., 2021). They facilitate carbon translocation to the host at lower rates and support lower coral growth rates (Pettay et al., 2015), which is all reflected in the physiological baseline changes observed in our branching corals. Such a shift to a thermotolerant symbiont community is another mechanism that may potentially underlie the reduced photosynthetic efficiency observed in the branching corals after thermal preconditioning. However, this is deemed less likely to occur in all the different distantly related coral species and within the short time frame of thermal preconditioning applied in this experiment (Boulotte et al., 2016; Scharfenstein et al., 2022).

### Thermal preconditioning enhances coral heat-stress tolerance

Our aquarium experiments demonstrated that thermal preconditioning can be applied to stress-harden corals and enhance their ability to cope with an acute heat stress pulse, contributing to the growing body of literature that has primarily reported this phenomenon from the field, including only few *ex-situ* studies (Ainsworth et al., 2016; Bay & Palumbi, 2015; Brown et al., 2023; Hackerott et al., 2021; Majerova et al., 2021; Martell, 2023; Wall et al., 2023). At the same time, our systematic and multi-species comparison has revealed that the receptiveness to the stress-hardening effects of thermal preconditioning strongly varied between the coral species. Overall, massive-growing species were most receptive to the stress-hardening effect of the thermal preconditioning treatments. In some species, we recorded increases of over 80 % in stress tolerance, while in others, increases were minor to nonexistent. This inter-species variation of stress-hardening receptiveness may explain the partially conflicting results across studies, considering that the studies that produced equivocal results also used different coral species in their investigations (Barshis et al., 2018; Henley et al., 2022; Klepac & Barshis, 2020; Schoepf et al., 2022). To explain these inter-species differences, we should, on the one hand, consider the thermal priming conditions, such as the mean exposure temperature and the diel temperature amplitude in relation to the upper thermal tolerance limit of a respective coral (Darling et al., 2012; Rivest et al., 2017). On the other hand, the individual acclimatization capacity and/or life-history strategy of the coral can be decisive for the receptiveness (Darling et al., 2012; Dilworth et al., 2021; Kenkel et al., 2013; Kenkel & Matz, 2016). For instance, some coral species might be unable to expand their thermal tolerance range beyond their absolute biological limits regardless of exposure to a priming stimulus (Klepac & Barshis, 2020; Schoepf et al., 2015, 2021).

Based on our data, we conclude that the inherent thermal tolerance of a species was not a good predictor for stress-hardening receptiveness. For example, species with high inherent thermal tolerance, such as *M. digitata*, were less receptive than other species with high inherent thermal tolerance (*G. fascicularis*) or thermally sensitive species, such as *A. muricata* and *P. verrucosa* (Fig. 6). This is in line with other studies showing that a stress-tolerant *Porites lobata* when acclimatized to a highly variable thermal environment became sensitive to bleaching, whereas a thermally susceptible *Acropora* sp. increased its bleaching tolerance after experiencing a similar high-variability condition (Klepac & Barshis, 2020; Thomas et al., 2018). It appears that the receptiveness to thermal preconditioning is likely connected to the priming stimulus and unique thermal tolerance range and life-history traits of each respective coral species. Hence, an empirical assessment of individual species-specific stress-hardening receptiveness is recommended prior to devising larger scale preconditioning efforts.

**Fig. 6.**
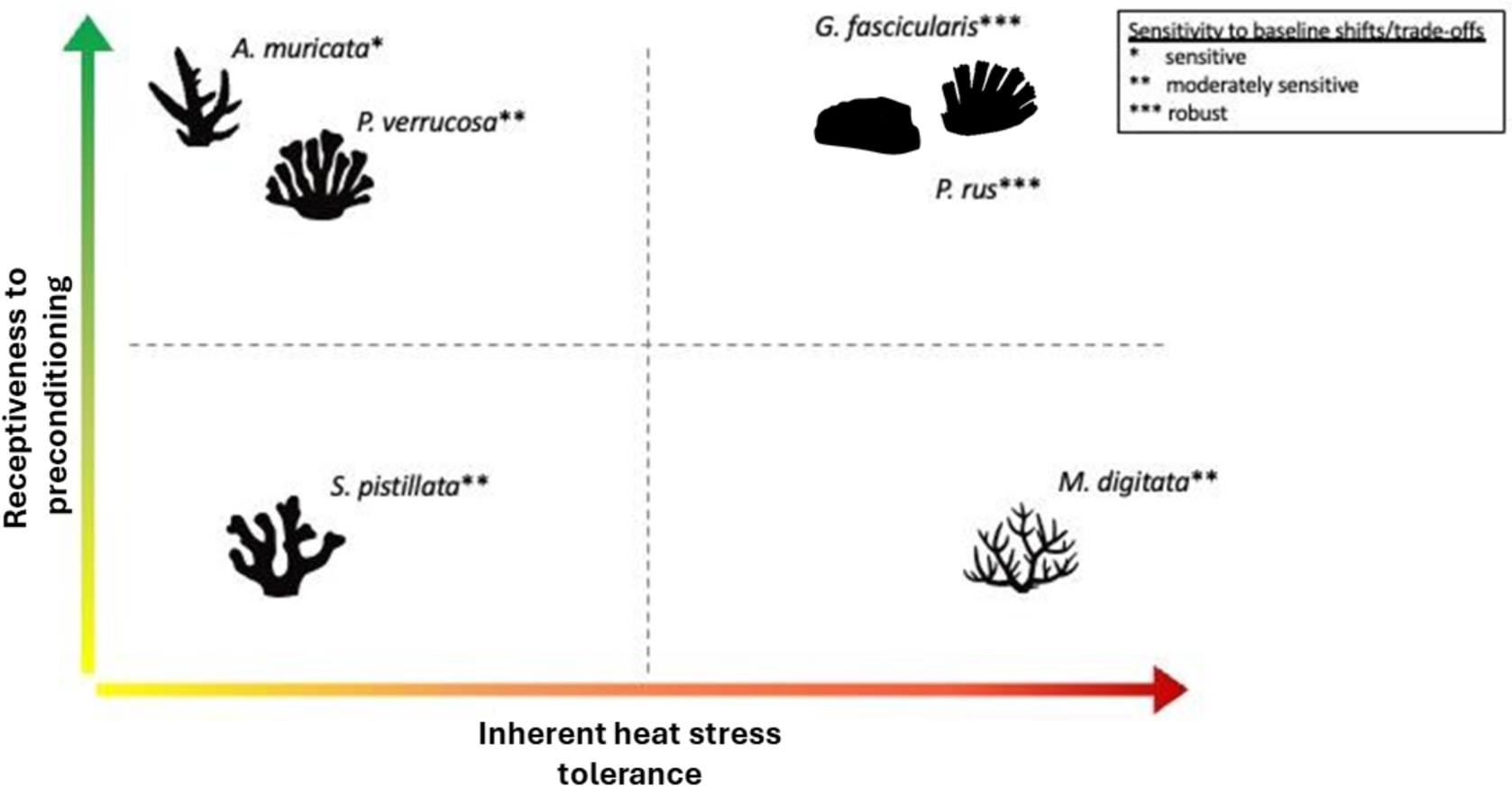
Conceptual representation of thermal stress-hardening receptiveness vs. inherent thermal tolerance and susceptibility to baseline shifts upon preconditioning in six coral species. The visual representation of each species’ receptiveness to priming as it is related to their inherent thermal tolerance provides valuable insights to help customize stress-hardening protocols for these species. Corals demonstrating high values in both metrics are excellent candidates for restoration programs that aim to use stress-hardening protocols, as these species can withstand extreme thermal anomalies and further increase their thermal tolerance through exposure to stress-hardening regimes. Highly receptive species with low inherent thermal tolerance are also promising for such programs, although local environmental conditions will require careful consideration, especially in heat-wave-prone areas. Species with high inherent thermal tolerance but limited receptiveness to priming may be suitable for restoration programs without priming, as stress-hardening treatments do not lead to further gains in thermal tolerance. Conversely, species in the lower-left corner, characterized by sensitivity to environmental stressors and low receptiveness to priming, may not be ideal candidates for such restoration programs. For these species, alternative stress-hardening approaches are suggested.

### Thermal preconditioning enhances resilience

Our study demonstrated that corals, previously exposed to thermal stress-hardening regimes, had higher survival and physiological recovery rates 30 days after acute heat stress exposure. Interestingly, the positive effects of the VT treatment excelled over the ST treatment. Preconditioned *P. rus* and *G. fascicularis* from the VT treatment fully recovered their physiological parameters after 30 days, indicating higher resilience than corals from the ST treatment, where physiological metrics did not fully recover the losses after heat stress. Similarly, other laboratory-based experiments and *in-situ* studies have shown that resilience was superior in corals from high-variability environments compared to other habitats (Oliver & Palumbi, 2011; Padilla-Gamiño et al., 2024; Schoepf et al., 2015; Wall et al., 2023) suggesting that fluctuating temperatures promote metabolic flexibility, allowing corals to optimize energy use and the repair mechanisms to recover better and faster.

We show that the inherent thermal tolerance of corals was a predictor for their long-term recovery trajectory. As such, species with inherently higher heat stress tolerance (*G. fascicularis, P.rus, M. digitata*) became more resilient and recovered at higher rates after thermal preconditioning and *vice versa,* thermally sensitive corals did not survive and recover well (*A. muricata*) despite the initial stress mitigation achieved through preconditioning. One explanation for the lower resilience in the inherently thermally sensitive species may be that the heat stress assay exceeded their upper thermal limit, beyond which preconditioning treatments could not further rescue the corals. This align with observations in the reef where species from the *Acroporidae* and *Pocilloporidae* families are known to suffer high mortalities during bleaching events and have a weak capacity to recover (Baird & Marshall, 2002; Burt et al., 2011). Accordingly, in our experiment, the low recovery and high mortality of *A. muricata*, *P. verrucosa*, and *S. pistillata,* might be linked to their competitive life-history strategy and lower upper thermal limits (Darling et al., 2012; McClanahan et al., 2007). Nevertheless, *Acropora*-dominated intertidal reefs also bleached less and recovered faster than communities from more thermally stable subtidal habitats (Camp et al., 2017; Le Nohaïc et al., 2017; Padilla-Gamiño et al., 2024). Reconciling these observations, we conclude that preconditioning treatments, when applied in appropriate doses, have the potential to enhance the resilience of these sensitive (branching) coral species and can help maintain stability of their communities.

During the recovery phase, the higher symbiont performance of ST and VT groups was linked to a significant increase in the survival and recovery rate, suggesting that a limited number of well-performing symbionts may be enough to support coral recovery (Fitt et al., 2001; Middlebrook et al., 2012). Our findings further confirm the work by Middlebrook et al. (2012), who showed that corals exposed to priming stimuli exhibited enhanced photosynthetic efficiency during heat stress, while the susceptibility to bleaching remained unchanged. This finding suggests that in stress-hardened corals, symbiont performance may be more important than symbiont density in determining coral resilience (Fitt et al., 2001; Martell, 2023; Middlebrook et al., 2012).

### Identifying the optimal thermal regime to efficiently stress harden corals

#### Thermal ranges and limits

The features of thermal regimes, including temperature increase, the fluctuation amplitude, temperature maxima and minima, and exposure duration, will be crucial to the success of the priming stimulus (Brown et al., 2023; Brown & Barott, 2022; Hackerott et al., 2021; Martell, 2023; Middlebrook et al., 2008). In our study, both preconditioning regimes, ST and VT, effectively induced various stress-hardening effects, including increases in stress tolerance and/or resilience, in the six coral species applying a mean temperature of 29 °C, i.e. 3 °C above the ambient mean temperature. Treatments were designed to remain within the previously identified 31 °C long-term upper limit of the corals inside the facility (Reichert et al., 2021). While Majerová & Drury (2022) also successfully documented stress-hardening effects by exposing corals to 29 °C (also 3 °C above the ambient control temperature), a temperature increase of only 2 °C above the ambient control temperature may be insufficient to trigger stress-hardening effects as shown in Henley et al. (2022). Similarly, priming corals with more extreme daily temperature fluctuations that exceed the range of 3 °C might already be overly stressful for the corals, and can hence explain the lack of stress-hardening effects or the reports of detrimental effects (Schoepf et al., 2021).

#### Thermal stability vs variability

Temperatures in the VT treatment underwent diel fluctuations with an amplitude of 3 °C from 27.5 to 30.5 °C. Despite the same mean temperatures applied in the two preconditioning treatments, the thermal tolerance of corals preconditioned with the VT treatment was increased by approximately 10 % compared to corals from the ST treatment. This is in line with the large body of literature showing that organisms from variable habitats are more stress tolerant than those from stable habitats (Bay & Palumbi, 2015; Drury et al., 2022; Oliver & Palumbi, 2011; Padilla-Gamiño et al., 2024), including specifically for corals from habitats with diel thermal variability of 2 to 3 °C (Brown et al., 2023). Moreover, temperature fluctuations have been shown to mitigate the responses to additional stressors, such as heavy metals in freshwater organisms, indicating a general improvement in stress mitigation and detoxification mechanisms (Hallman & Brooks, 2015). In these studies, not only temperature maxima but also the minima likely determined the priming outcomes (Drury et al., 2022; Hallman & Brooks, 2015; Schoepf et al., 2019; Wall et al., 2023). For instance, an *in situ* study reported that exposure to cooler temperature pulses alone, i.e., fluctuation without any extremely high-temperature peaks, can successfully induce stress-hardening in corals and may provide a safer option compared to the implementation of high-temperature pulses (Wall et al., 2023). Overall, the similar outcomes of the ST and VT preconditioning treatments in our study suggests that the stress-hardening effect, in a large part, is stimulated by the increase of the ambient temperature by 3 °C. Yet, implementing thermal variability rather than a stable high-temperature regime as a stimulus for the preconditioning of corals promises to induce stress hardening more effectively, by up to 10 % based on our results.

#### Species receptiveness to thermal preconditioning

Our study demonstrated that coral species differ in receptiveness to stress-hardening treatments. We classified the coral species *G. fascicularis, P. rus, and M. digitata* species as thermally resistant and *A. muricata, P. verrucosa,* and *S. pistillata* as thermally sensitive. Among these, *G. fascicularis, A. muricata, P. rus,* and *P. verrucosa* were receptive to stress-hardening, whereas *M. digitata* and *S. pistillata* were non-receptive. For future investigations and practitioner action, we provide a simplified conceptual classification of the stress-hardening receptiveness of coral species in relation to their inherent thermal tolerance and their susceptibility to baseline physiological shifts, with potential consequences (Fig. 6). The chart provides an index for optimizing restoration efforts, ensuring resource effectiveness, and can help interpret and better understand outcomes across stress-hardening studies, helping to uncover complex mechanisms that may underlie the equivocal results from the yet non-standardized studies.

## Conclusions

This study examined the effects of three thermal preconditioning regimes on the heat stress tolerance, recovery, and survival of six stony coral species. We showed that thermal preconditioning using a variable-high temperature was most effective in increasing thermal tolerance and promoting recovery after an acute heat stress. However, the stress-hardening receptiveness of coral species differed and physiological shifts after preconditioning occurred, which may entail undesirable trade-offs in the long term, deeming some coral species unsuitable for thermal stress-hardening applications in restoration programs. While the receptiveness to stress-hardening treatments and the subsequent increase in thermal tolerance was independent of the inherent thermal tolerance of the species, the boost in survival and recovery capacity attained through preconditioning was greater in corals characterized by higher inherent thermal tolerance. A deeper understanding of the underlying physiological and metabolic mechanisms that act in thermal preconditioning of corals and identifying thermal stress-hardening regimes that maximize thermal tolerance without exposing species to harmful trade-offs will be crucial.

## Acknowledgments

We thank Patrick Schubert and Christina Anding from the Justus Liebig University Giessen, Germany for their support in animal caretaking and maintenance of the research facilities as well as the Marine Holobiomics Lab for the everyday background support. We also thank Miriam Altvatter, Antonio Pinto, Mariangela Impalà, André Dietzmann, Philipp Seibold for their assistance and for their contribution during experiments.

## Author contributions

E.F.F., A.R., and M.Z. conceived the study and designed the experiments E.F.F. and F.W-Z collected data, E.F.F. analyzed and curated data. E.F.F., A.R., and M.Z. wrote and revised the manuscript. M.Z. provided research materials and logistics. All authors read and approved the manuscript.

## Availability of data and materials

Additional data, analyses and visualization are presented in the Supplementary materials. All original data and analysis code are accessible via the *GitHub* repository under the accession link: https://github.com/ErikFerrara/Stress-hardening.git

## Declarations

The authors declare that they have no competing interests.

## Funding

E.F.F. was supported by a postgraduate stipend of Justus Liebig University Giessen. The project was conducted in the ‘Ocean 2100’ facilities of the Justus Liebig University Giessen, which is part of the global change simulation project of the Colombian-German Center of Excellence in Marine Sciences (CEMarin). AR acknowledges the funding of the Helmholtz Institute for Functional Marine Biodiversity at the University of Oldenburg, Niedersachsen, Germany. HIFMB is a collaboration between the Alfred-Wegener Institute, Helmholtz Center for Polar and Marine Research, and the Carl-von-Ossietzky University Oldenburg. It was initially funded by the Ministry for Science and Culture of Lower Saxony and the Volkswagen Foundation through the “Niedersächsisches Vorab” grant program (grant number ZN3285).

## Supplementary materials

### Supplementary Tables

**Table S1:**
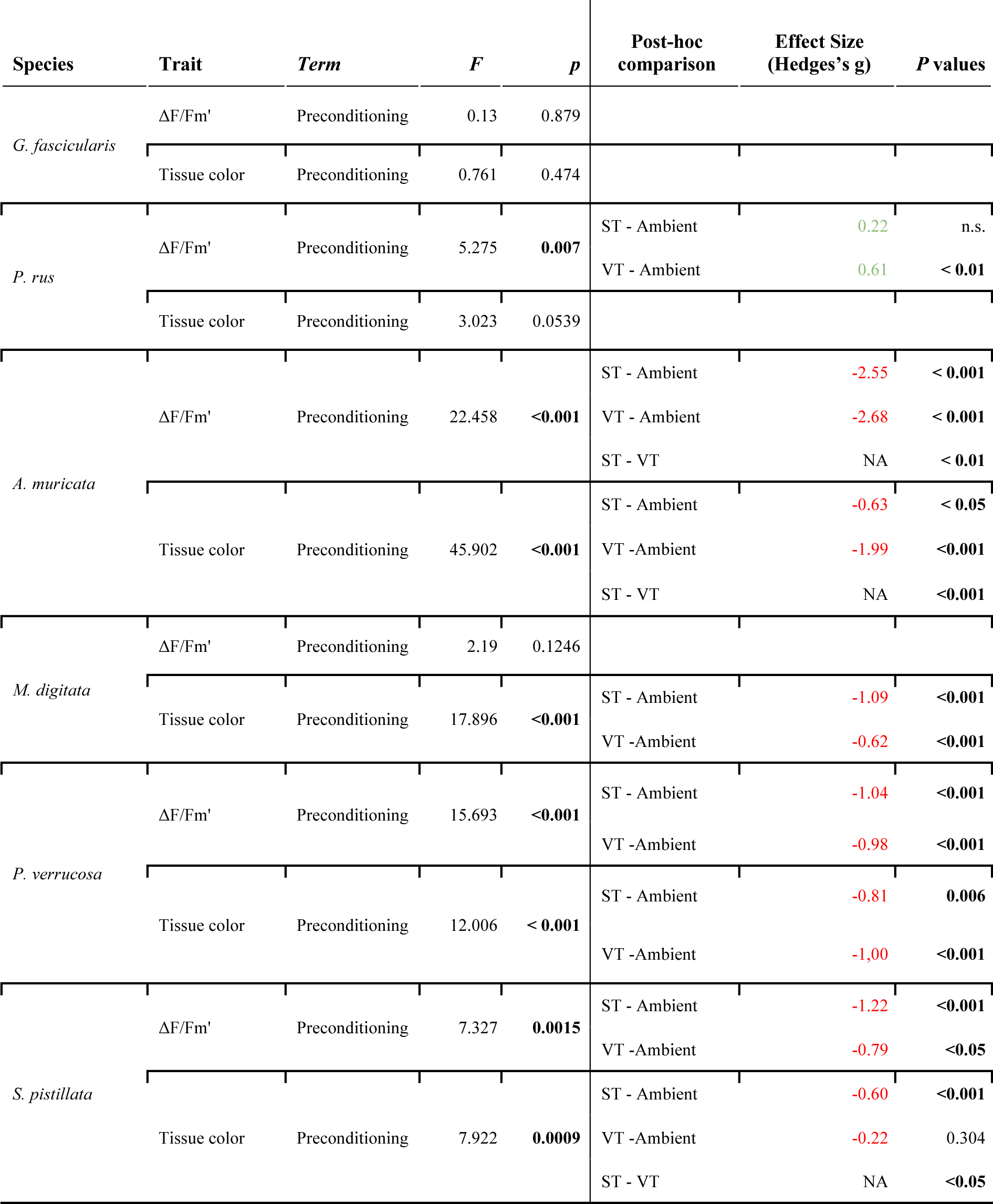
Statistical results of thermal preconditioning regime on baseline coral physiology. The F statistic was calculated using the ANOVA test on the liner mixed effects model. Effect size, shown as hedges g, was estimated as the difference between treatments comparing the raw values of the stable-Ambient group with the two preconditioning treatments.

**Table S2:** Effective quantum yield (ΔF/Fm’), median delta (post-heat minus post-preconditioning) and percentage reduction in heat stress response (e.g. ΔST minus ΔAmbient).

| Species | $\Delta Ambient$ | $\Delta ST$ | $\Delta VT$ | $\Delta ST$ minus<br>$\Delta Ambient$ (%) | $\Delta VT$ minus<br>$\Delta Ambient$ (%) | $VT$ minus<br>$ST$ (%) |
| --- | --- | --- | --- | --- | --- | --- |
| *G. fascicularis* | -0.128 | -0.012 | -0.015 | **90.66** | 88.72 | -1.95 |
| *P. rus* | -0.459 | -0.255 | -0.195 | 44.34 | **57.41** | **+13.07** |
| *A. muricata* | -0.378 | -0.078 | -0.069 | 79.52 | **81.64** | **+2.11** |
| *M. digitata* | -0.071 | -0.047 | -0.028 | 33.57 | **60.14** | **+26.57** |
| *P. verrucosa* | -0.458 | -0.253 | -0.188 | 44.76 | **58.84** | **+14.08** |
| *S. pistillata* | -0.551 | -0.408 | -0.382 | 26.04 | **30.67** | **+4.63** |

**Table S3:** Median delta of tissue color intensity (post-heat minus post-preconditioning) and percentage reduction in heat stress response (e.g. ΔST minus ΔAmbient).

| Species | ΔAmbient | ΔST | ΔVT | ΔST minus<br>ΔAmbient (%) | ΔVT minus<br>ΔAmbient (%) | VT minus<br>ST (%) |
| --- | --- | --- | --- | --- | --- | --- |
| *G. fascicularis* | -60.973 | -28.178 | -26.610 | 53.79 | **56.36** | **+2.57** |
| *P. rus* | -38.454 | -27.802 | -31.386 | **27.70** | 18.38 | -9.32 |
| *A. muricata* | -128.826 | -98.456 | -86.824 | 23.58 | **32.60** | **+9.03** |
| *M. digitata* | -43.772 | -22.870 | -25.168 | **47.75** | 42.50 | -5.25 |
| *P. verrucosa* | -87.950 | -69.782 | -61.224 | 20.66 | **30.39** | **+9.73** |
| *S. pistillata* | -103.238 | -96.879 | -98.204 | **6.16** | 4.88 | -1.28 |

**Table S4:** Statistical results of the response of preconditioned corals to heat stress (post-preconditioning vs. heat). Significant p-values (Wilcoxon tests) are shown in bold. Effect size, shown as Hedges g, was estimated within each treatment by comparing post-preconditioning minus post-heat paired values.

| Species | Analysis | Term | Contrast<br>(Post-heat - Post-preconditioning) | P value<br>(Wilcoxon test) | Signif. | Effect<br>size |
| --- | --- | --- | --- | --- | --- | --- |
| *G. fascicularis* | $\Delta F/Fm'$ | Preconditioning | Ambient | **<0.001** | *** | -1.70 |
|  |  |  | ST | **0.01** | ** | -1.35 |
|  |  |  | VT | 0.103 |  | -0.85 |
|  | Tissue color | Preconditioning | Ambient | **<0.001** | *** | -2.43 |
|  |  |  | ST | **<0.001** | *** | -3.42 |
|  |  |  | VT | **0.001** | *** | -2.32 |
| *P. rus* | $\Delta F/Fm'$ | Preconditioning | Ambient | **<0.001** | *** | -3.08 |
|  |  |  | ST | **<0.001** | *** | -1.77 |
|  |  |  | VT | **<0.001** | *** | -1.76 |
|  | Tissue color | Preconditioning | Ambient | **<0.001** | *** | -2.72 |
|  |  |  | ST | **<0.001** | *** | -1.28 |
|  |  |  | VT | **<0.001** | *** | -1.65 |
| *A. muricata* | $\Delta F/Fm'$ | Preconditioning | Ambient | **<0.001** | *** | -4.35 |
|  |  |  | ST | **<0.01** | ** | -1.16 |
|  |  |  | VT | **<0.05** | * | -0.9 |
|  | Tissue color | Preconditioning | Ambient | **<0.001** | *** | -9.88 |
|  |  |  | ST | **<0.001** | *** | -8.06 |
|  |  |  | VT | **<0.001** | *** | -4.54 |
| *M. digitata* | $\Delta F/Fm'$ | Preconditioning | Ambient | **0.004** | ** | -0.99 |
|  |  |  | ST | **0.003** | ** | -1.95 |
|  |  |  | VT | **0.021** | * | -1.39 |
|  | Tissue color | Preconditioning | Ambient | **0.003** | ** | -1.95 |
|  |  |  | ST | **0.021** | * | -1.25 |
|  |  |  | VT | **0.038** | * | -1.18 |
| *P. verrucosa* | $\Delta F/Fm'$ | Preconditioning | Ambient | **<0.001** | *** | -4.3 |
|  |  |  | ST | **<0.001** | *** | -1.92 |
|  |  |  | VT | **<0.001** | *** | -2.35 |
|  | Tissue colo | Preconditioning | Ambient | **<0.001** | *** | -6.05 |
|  |  |  | ST | **<0.001** | *** | -4.09 |
|  |  |  | VT | **<0.001** | *** | -3.78 |
| *S. pistillata* | $\Delta F/Fm'$ | Preconditioning | Ambient | **<0.001** | *** | -17.08 |
|  |  |  | ST | **<0.001** | *** | -4.05 |
|  |  |  | VT | **<0.001** | *** | -4.9 |
|  | Tissue color | Preconditioning | Ambient | **<0.001** | *** | -5.42 |
|  |  |  | ST | **<0.001** | *** | -5.86 |
|  |  |  | VT | **<0.001** | *** | -5.76 |

**Table S5:** Statistical comparison of coral receptiveness between preconditioning treatments based on paired delta values (Δ=post-heat minus post-preconditioning). Significant Bonferroni adjusted p-values (linear mixed effect model) are highlighted in bold.

| Species | Analysis ($\Delta$ values) | Term | Contrast<br>(Pairwais post-hoc) | *P* value<br>(Adj. BH) | Signif. |
| --- | --- | --- | --- | --- | --- |
| *G. fascicularis* | $\Delta F/Fm'$ | Preconditioning | ΔAmbient - ΔST | **<0.01** | ** |
| | | | ΔAmbient - ΔVT | **<0.001** | *** |
| | Tissue color | Preconditioning | ΔAmbient - ΔST | **<0.001** | *** |
| | | | ΔAmbient - ΔVT | **<0.001** | *** |
| *P. rus* | $\Delta F/Fm'$ | Preconditioning | ΔAmbient - ΔST | **<0.01** | ** |
| | | | ΔAmbient - ΔVT | **<0.001** | *** |
| | Tissue color | Preconditioning | ΔAmbient - ΔST | 0.0921 | |
| | | | ΔAmbient - ΔVT | 0.201 | |
| *A. muricata* | $\Delta F/Fm'$ | Preconditioning | ΔAmbient - ΔST | **<0.001** | *** |
| | | | ΔAmbient - ΔVT | **<0.001** | *** |
| | Tissue color | Preconditioning | ΔAmbient - ΔST | **<0.001** | *** |
| | | | ΔAmbient - ΔVT | **<0.001** | *** |
| | | | ΔST - ΔVT | **<0.05** | * |
| *M. digitata* | $\Delta F/Fm'$ | Preconditioning | ΔAmbient - ΔST | 1 | |
| | | | ΔAmbient - ΔVT | 0.968 | |
| | Tissue color | Preconditioning | ΔAmbient - ΔST | **<0.05** | * |
| | | | ΔAmbient - ΔVT | **<0.01** | ** |
| *P. verrucosa* | $\Delta F/Fm'$ | Preconditioning | ΔAmbient - ΔST | **<0.001** | *** |
| | | | ΔAmbient - ΔVT | **<0.001** | *** |
| | Tissue color | Preconditioning | ΔAmbient - ΔST | **<0.01** | ** |
| | | | ΔAmbient - ΔVT | **<0.001** | *** |
| *S. pistillata* | $\Delta F/Fm'$ | Preconditioning | ΔAmbient - ΔST | **<0.001** | *** |
| | | | ΔAmbient - ΔVT | **<0.001** | *** |
| | Tissue color | Preconditioning | ΔAmbient - ΔST | 1 | |
| | | | ΔAmbient - ΔVT | 1 | |

**Table S6:** Statistical results of recovery of preconditioned corals after 30 days. Recovery was estimated as the difference (effect size as hedges g) between the heat and control treatments within each treatment for each response variable and across species. Significant p-values (two-tailed permutation t-test) are shown in bold. Missing measurements due to death of all fragments are indicated by “NA”.

| Species | Analysis | Term | Contrast<br>(heat - control) | Effect size<br>(hedges_g) | P value<br>(two-sided permutation t-test) | Signif. |
| --- | --- | --- | --- | --- | --- | --- |
| *G. fascicularis* | $\Delta F/Fm'$<br>(recovery) | Preconditioning | Ambient | -1.20 | **0.028** | * |
|  |  |  | ST | -2.55 | **0.028** | * |
|  |  |  | VT | -1.37 | 0.114 |  |
|  | Tissue color<br>(recovery) | Preconditioning | Ambient | -1.37 | **0.028** | ** |
|  |  |  | ST | -0.95 | 0.2 |  |
|  |  |  | VT | -0.98 | 0.114 |  |
| *P. rus* | $\Delta F/Fm'$<br>(recovery) | Preconditioning | Ambient | -1.20 | **<0.001** | *** |
|  |  |  | ST | -2.55 | **0.015** | * |
|  |  |  | VT | -1.37 | 0.720 |  |
|  | Tissue color<br>(recovery) | Preconditioning | Ambient | -2.72 | **<0.001** | *** |
|  |  |  | ST | -0.86 | **0.05** | * |
|  |  |  | VT | -0.70 | 0.065 |  |
| *A. muricata* | $\Delta F/Fm'$<br>(recovery) | Preconditioning | Ambient | NA | NA | |
|  |  |  | ST | NA | NA |  |
|  |  |  | VT | NA | NA |  |
|  | Tissue color<br>(recovery) | Preconditioning | Ambient | NA | NA |  |
|  |  |  | ST | NA | NA |  |
|  |  |  | VT | NA | NA |  |
| *M. digitata* | $\Delta F/Fm'$<br>(recovery) | Preconditioning | Ambient | -0.51 | 0.486 | |
|  |  |  | ST | -0.08 | 0.885 |  |
|  |  |  | VT | -0.91 | 0.114 |  |
|  | Tissue color<br>(recovery) | Preconditioning | Ambient | -0.37 | 0.342 |  |
|  |  |  | ST | -0.28 | 1 |  |
|  |  |  | VT | 0.25 | 0.685 |  |
| *P. verrucosa* | $\Delta F/Fm'$<br>(recovery) | Preconditioning | Ambient | NA | NA | |
|  |  |  | ST | NA | NA |  |
|  |  |  | VT | NA | NA |  |
|  | Tissue color<br>(recovery) | Preconditioning | Ambient | NA | NA |  |
|  |  |  | ST | NA | NA |  |
|  |  |  | VT | NA | NA |  |
| *S. pistillata* | $\Delta F/Fm'$<br>(recovery) | Preconditioning | Ambient | NA | NA | |
|  |  |  | ST | NA | NA |  |
|  |  |  | VT | NA | NA |  |
|  | Tissue color<br>(recovery) | Preconditioning | Ambient | NA | NA |  |
|  |  |  | ST | NA | NA |  |
|  |  |  | VT | NA | NA |  |

**Table S7:**
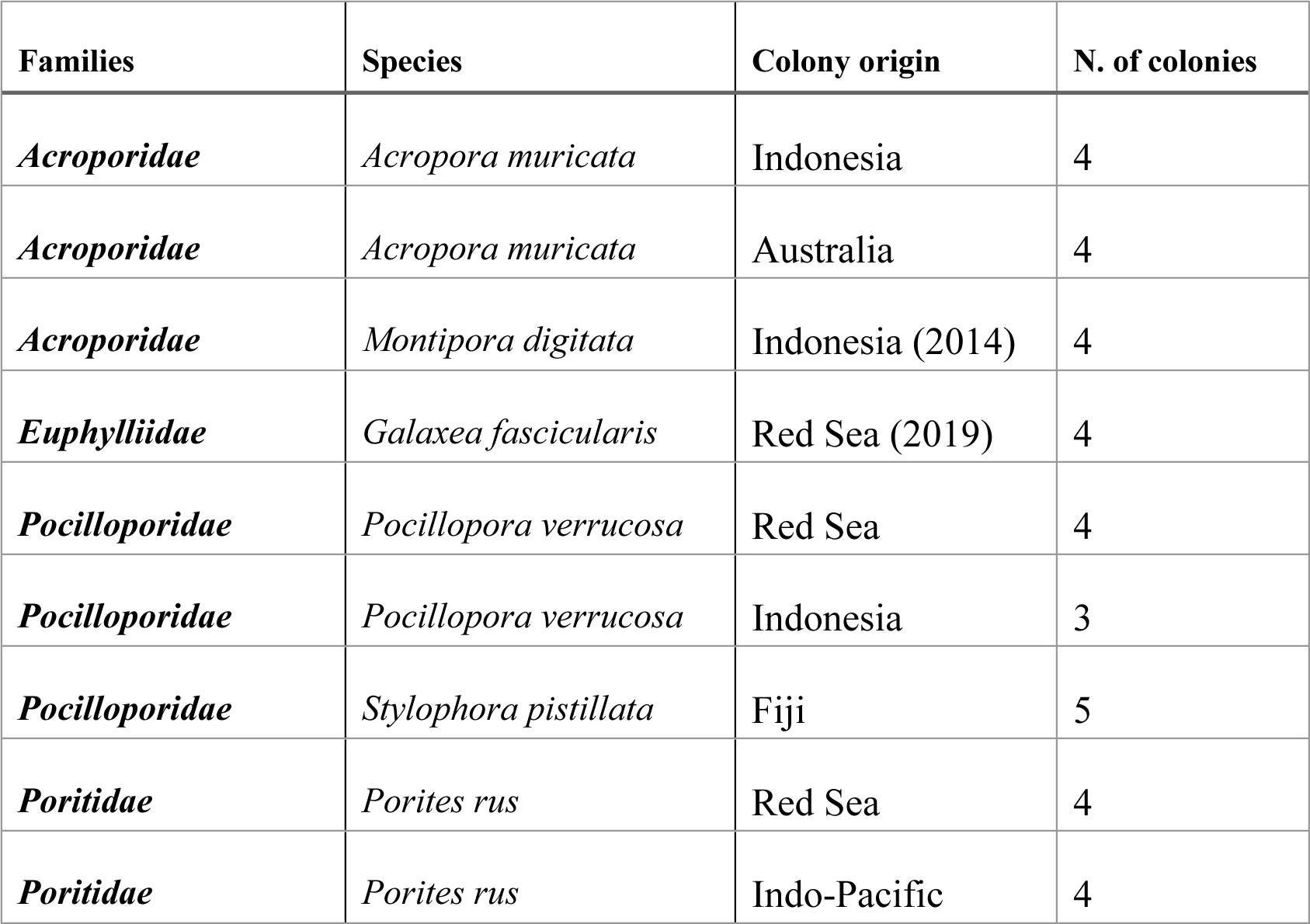
Coral species, countries of origin and number of colonies used during the experiment.

### Pilot study

The target temperatures for the heat stress assays to induce heat stress in the different coral species were determined in a pilot heat stress series. This pilot study aimed to identify the specific temperatures that would induce a severe but not lethal response in each of the coral species. This ideal heat stress temperature can differ between species of different heat tolerances. To this end, test fragments from each species were exposed to 34 °C, 35 °C, or 36 °C following two established protocols (Doering et al., 2021; Reichert et al., 2021). We observed a measurable increase in the heat stress responses between 34° and 35°C, while 36°C was lethal for most species. Based on the observations in this pilot study, we selected the target temperature of 35 °C, i.e., 9 °C above the ambient mean temperature of the facility.

### Supplementary Figures

**Figure S1.**
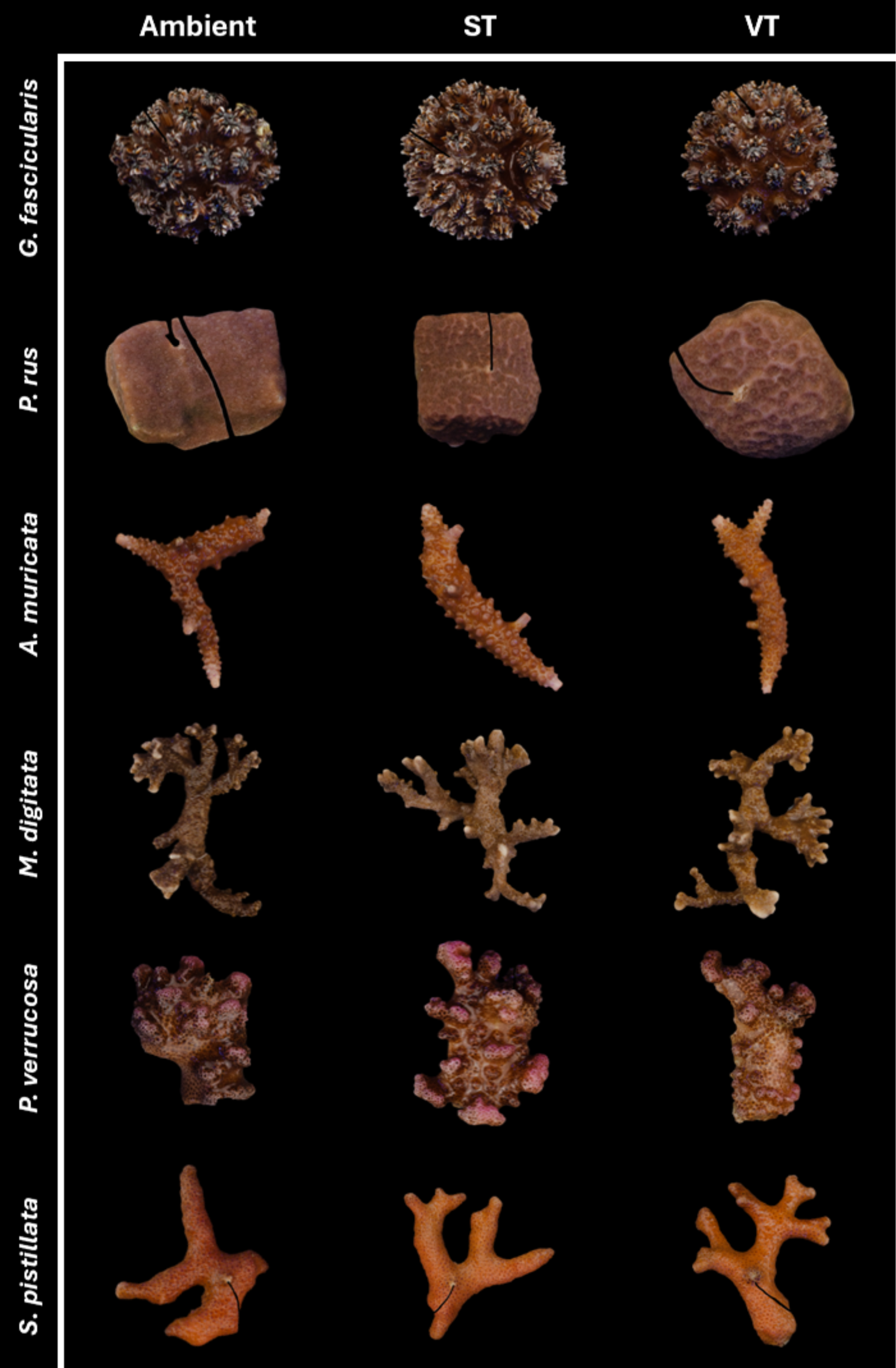
Tissue color changes after preconditioning before the heat stress assays. Comparison of coral tissue color between preconditioning treatment at the “Post-preconditioning” time point. Some of the corals from ST and VT treatments were slightly paler than the Ambient as a result of the stress-hardening process.

**Figure S2.**
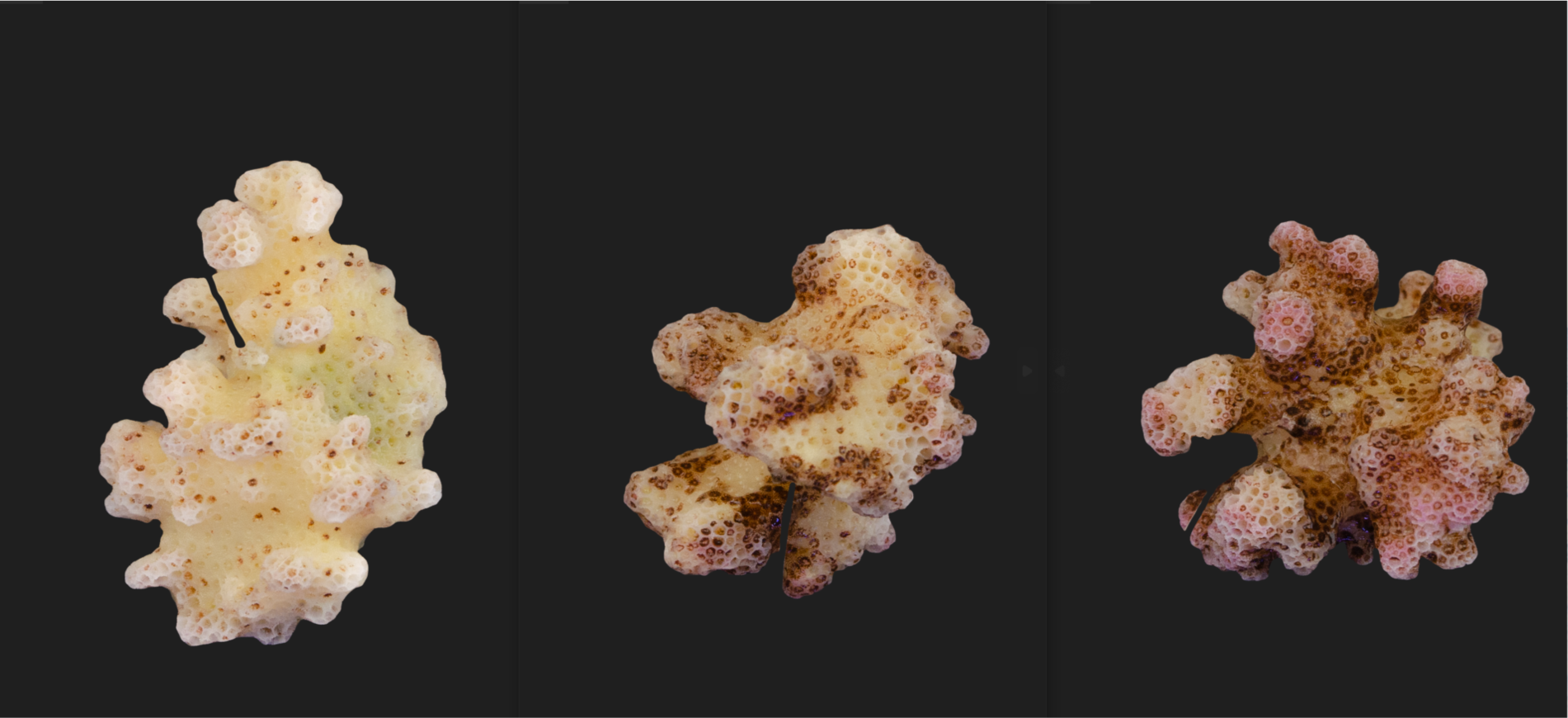
*Pocillopora verrucosa* responses to heat stress. The picture shows the response to the heat of the same colony preconditioned with distinct thermal regimes. On the left, the colony was kept at ambient temperature (26°C) before the heat stress. In the middle, the colony was preconditioned with the stable-high temperature regime, while on the right, with the variable-high temperature.

**Figure S3.**
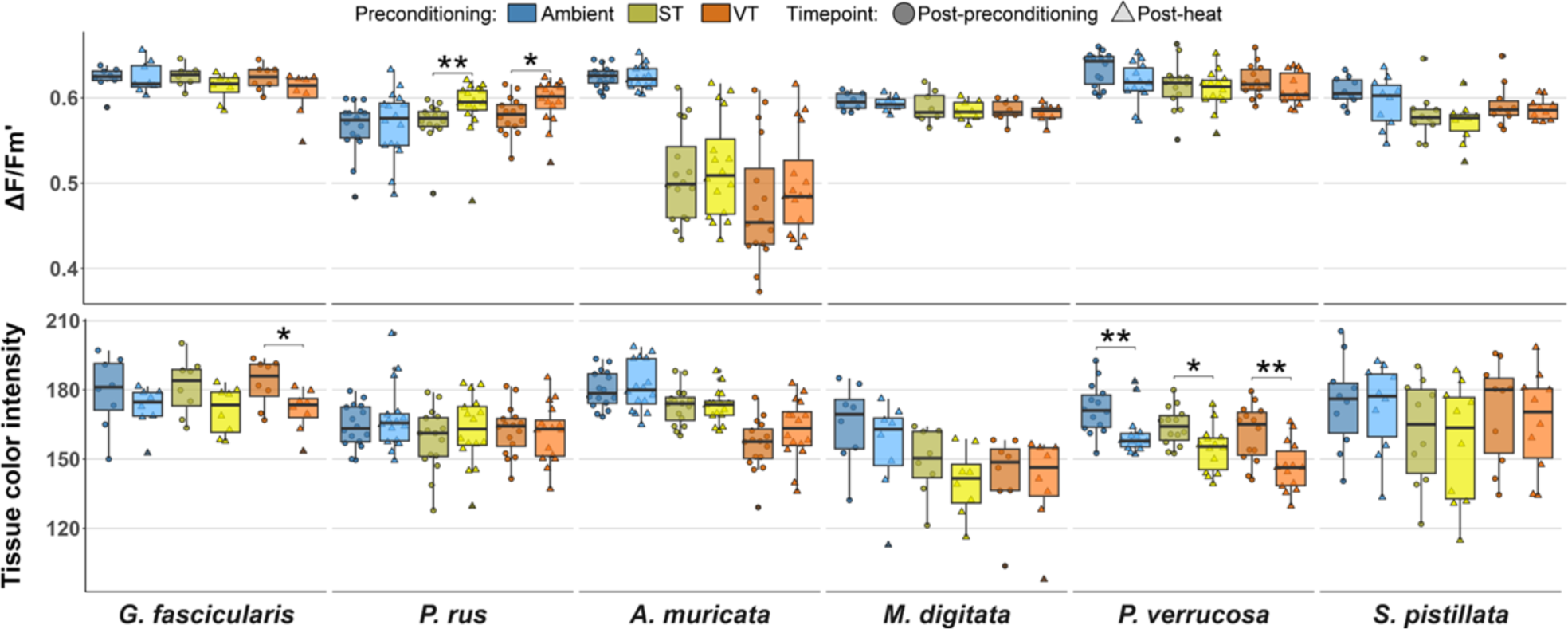
Post-heat paired Control. The changes in effective quantum yield (ΔF/Fm’) (A) and tissue color intensity (C) within each preconditioning group of the “Control” treatments are shown as boxplots, comparing paired “post-heat” (lighter color) to “post-preconditioning” (darker color) values. Data are displayed as boxplots with raw data points. Connecting lines between boxes indicate significant differences between time points (p < 0.001***, < 0.01**, < 0.05* from Kruskal-Wallis and post hoc Wilcoxon test).

## Notes

### Competing Interest Statement

The authors have declared no competing interest.

## References

Ainsworth, T. D., Heron, S. F., Ortiz, J. C., Mumby, P. J., Grech, A., Ogawa, D., Eakin, C. M., & Leggat, W. (2016). Climate change disables coral bleaching protection on the Great Barrier Reef. Science, 352(6283), 338–342. 10.1126/science.aac7125

Baird, A. H., & Marshall, P. A. (2002). Mortality, growth, and reproduction in scleractinian corals following bleaching on the Great Barrier Reef. Marine Ecology Progress Series, 237, 133–141. 10.3354/meps237133

Barshis, D. J., Birkeland, C., Toonen, R. J., Gates, R. D., & Stillman, J. H. (2018). High-frequency temperature variability mirrors fixed differences in thermal limits of the massive coral Porites lobata. Journal of Experimental Biology, 221(24), jeb188581. 10.1242/jeb.188581

Bay, R. A., & Palumbi, S. R. (2015). Rapid acclimation ability mediated by transcriptome changes in reef-building corals. Genome Biology and Evolution, 7(6), 1602–1612. 10.1093/gbe/evv085

Bellantuono, A. J., Hoegh-Guldberg, O., & Rodriguez-Lanetty, M. (2011). Resistance to thermal stress in corals without changes in symbiont composition. Proceedings of the Royal Society B: Biological Sciences, 279(1731), 1100–1107. 10.1098/rspb.2011.1780

Bellantuono, A. J., Hoegh-Guldberg, O., & Rodriguez-Lanetty, M. (2012). Resistance to thermal stress in corals without changes in symbiont composition. Proceedings of the Royal Society B: Biological Sciences, 279(1731), 1100–1107. 10.1098/rspb.2011.1780

Brown, K., & Barott, K. L. (2022). The Costs and Benefits of Environmental Memory for Reef-Building Corals Coping with Recurring Marine Heatwaves. Integrative and Comparative Biology, 62(6), 1748–1755. 10.1093/icb/icac074

Brown, K., Martynek, M., & Barott, K. L. (2023). Maximal coral thermal tolerance is found at intermediate diel temperature variability (p. 2023.03.27.534434). bioRxiv. 10.1101/2023.03.27.534434

Burt, J., Al-Harthi, S., & Al-Cibahy, A. (2011). Long-term impacts of coral bleaching events on the world’s warmest reefs. Marine Environmental Research, 72(4), 225–229. 10.1016/j.marenvres.2011.08.005

Calabrese, E. J., Bachmann, K. A., Bailer, A. J., Bolger, P. M., Borak, J., Cai, L., Cedergreen, N., Cherian, M. G., Chiueh, C. C., Clarkson, T. W., Cook, R. R., Diamond, D. M., Doolittle, D. J., Dorato, M. A., Duke, S. O., Feinendegen, L., Gardner, D. E., Hart, R. W., Hastings, K. L., … Mattson, M. P. (2007). Biological stress response terminology: Integrating the concepts of adaptive response and preconditioning stress within a hormetic dose–response framework. Toxicology and Applied Pharmacology, 222(1), 122–128. 10.1016/j.taap.2007.02.015

Calabrese, E. J., & Mattson, M. P. (2017). How does hormesis impact biology, toxicology, and medicine? Npj Aging and Mechanisms of Disease, 3(1), Articolo 1. 10.1038/s41514-017-0013-z

Camp, E. F., Nitschke, M. R., Rodolfo-Metalpa, R., Houlbreque, F., Gardner, S. G., Smith, D. J., Zampighi, M., & Suggett, D. J. (2017). Reef-building corals thrive within hot-acidified and deoxygenated waters. Scientific Reports, 7(1), Articolo 1. 10.1038/s41598-017-02383-y

Camp, E. F., Smith, D. J., Evenhuis, C., Enochs, I., Manzello, D., Woodcock, S., & Suggett, D. J. (2016). Acclimatization to high-variance habitats does not enhance physiological tolerance of two key Caribbean corals to future temperature and pH. Proceedings of the Royal Society B: Biological Sciences, 283(1831), 20160442. 10.1098/rspb.2016.0442

Cantin, N. E., van Oppen, M. J. H., Willis, B. L., Mieog, J. C., & Negri, A. P. (2009). Juvenile corals can acquire more carbon from high-performance algal symbionts. Coral Reefs, 28(2), 405–414. 10.1007/s00338-009-0478-8

Carelli, G., & Iavicoli, I. (2002). Defining hormesis: The necessary tool to clarify experimentally the low dose– response relationship. Human & Experimental Toxicology, 21(2), 103–104. 10.1191/0960327102ht219oa

Coles, S. L., & Brown, B. E. (2003). Coral bleaching—Capacity for acclimatization and adaptation. In Advances in Marine Biology (Vol. 46, pp. 183–223). Academic Press. 10.1016/S0065-2881(03)46004-5

Coles, S. L., & Jokiel, P. L. (1977). Effects of temperature on photosynthesis and respiration in hermatypic corals. Marine Biology, 43(3), 209–216. 10.1007/BF00402313

Cornwell, B., Armstrong, K., Walker, N. S., Lippert, M., Nestor, V., Golbuu, Y., & Palumbi, S. R. (2021). Widespread variation in heat tolerance and symbiont load are associated with growth tradeoffs in the coral Acropora hyacinthus in Palau. eLife, 10, e64790. 10.7554/eLife.64790

Cunning, R., & Baker, A. C. (2013). Excess algal symbionts increase the susceptibility of reef corals to bleaching. Nature Climate Change, 3. 10.1038/nclimate1711

Cunning, R., & Baker, A. C. (2014). Not just who, but how many: The importance of partner abundance in reef coral symbioses. Frontiers in Microbiology, 5. https://www.frontiersin.org/journals/microbiology/articles/10.3389/fmicb.2014.00400

Cunning, R., Silverstein, R. N., & Baker, A. C. (2018). Symbiont shuffling linked to differential photochemical dynamics of Symbiodinium in three Caribbean reef corals. Coral Reefs, 37(1), 145–152. 10.1007/s00338-017-1640-3

Darling, E. S., Alvarez-Filip, L., Oliver, T. A., McClanahan, T. R., & Côté, I. M. (2012). Evaluating life-history strategies of reef corals from species traits. Ecology Letters, 15(12), 1378–1386. 10.1111/j.1461-0248.2012.01861.x

DeMerlis, A., Kirkland, A., Kaufman, M. L., Mayfield, A. B., Formel, N., Kolodziej, G., Manzello, D. P., Lirman, D., Traylor-Knowles, N., & Enochs, I. C. (2022). Pre-exposure to a variable temperature treatment improves the response of Acropora cervicornis to acute thermal stress. Coral Reefs, 41(2), 435–445. 10.1007/s00338-022-02232-z

Drury, C. (2020). Resilience in reef-building corals: The ecological and evolutionary importance of the host response to thermal stress. Molecular Ecology, 29(3), 448–465. 10.1111/mec.15337

Drury, C., Dilworth, J., Majerová, E., Caruso, C., & Greer, J. B. (2022). Expression plasticity regulates intraspecific variation in the acclimatization potential of a reef-building coral. Nature Communications, 13(1), Articolo 1. 10.1038/s41467-022-32452-4

Ellegren, H., & Sheldon, B. C. (2008). Genetic basis of fitness differences in natural populations. Nature, 452(7184), Articolo 7184. 10.1038/nature06737

Evensen, N. R., Voolstra, C. R., Fine, M., Perna, G., Buitrago-López, C., Cárdenas, A., Banc-Prandi, G., Rowe, K., & Barshis, D. J. (2022). Empirically derived thermal thresholds of four coral species along the Red Sea using a portable and standardized experimental approach. Coral Reefs, 239–252. 10.1007/s00338-022-02233-y

Fine, M., & Loya, Y. (2002). Endolithic algae: An alternative source of photoassimilates during coral bleaching. Proceedings of the Royal Society of London. Series B: Biological Sciences, 269(1497), 1205–1210. 10.1098/rspb.2002.1983

Fitt, W., Brown, B., Warner, M., & Dunne, R. (2001). Coral bleaching: Interpretation of thermal tolerance limits and thermal thresholds in tropical corals. Coral Reefs, 20(1), 51–65. 10.1007/s003380100146

Fox, J., & Weisberg, S. (2019). R Companion 3E. https://www.john-fox.ca/Companion/index.html

Frölicher, T. L., Fischer, E. M., & Gruber, N. (2018). Marine heatwaves under global warming. Nature, 560(7718), Articolo 7718. 10.1038/s41586-018-0383-9

Gibbin, E. M., Krueger, T., Putnam, H. M., Barott, K. L., Bodin, J., Gates, R. D., & Meibom, A. (2018). Short-term thermal acclimation modifies the metabolic condition of the coral holobiont. Frontiers in Marine Science, 5(FEB), 1–11. 10.3389/fmars.2018.00010

Hackerott, S., Martell, H. A., & Eirin-Lopez, J. M. (2021). Coral environmental memory: Causes, mechanisms, and consequences for future reefs. Trends in Ecology and Evolution, 36(11), 1011–1023. 10.1016/j.tree.2021.06.014

Hallman, T. A., & Brooks, M. L. (2015). The deal with diel: Temperature fluctuations, asymmetrical warming, and ubiquitous metals contaminants. Environmental Pollution, 206, 88–94. 10.1016/j.envpol.2015.06.005

Helgoe, J., Davy, S. K., Weis, V. M., & Rodriguez-Lanetty, M. (2024). Triggers, cascades, and endpoints: Connecting the dots of coral bleaching mechanisms. Biological Reviews, n/a(n/a). 10.1111/brv.13042

Henley, E. M., Bouwmeester, J., Jury, C. P., Toonen, R. J., Quinn, M., Lager, C. V. A., & Hagedorn, M. (2022). Growth and survival among Hawaiian corals outplanted from tanks to an ocean nursery are driven by individual genotype and species differences rather than preconditioning to thermal stress. PeerJ. 10.7717/peerj.13112

Hilker, M., Schwachtje, J., Baier, M., Balazadeh, S., Bäurle, I., Geiselhardt, S., Hincha, D. K., Kunze, R., Mueller-Roeber, B., Rillig, M. C., Rolff, J., Romeis, T., Schmülling, T., Steppuhn, A., van Dongen, J., Whitcomb, S. J., Wurst, S., Zuther, E., & Kopka, J. (2016). Priming and memory of stress responses in organisms lacking a nervous system. Biological Reviews, 91(4), 1118–1133. 10.1111/brv.12215

Ho, J., Tumkaya, T., Aryal, S., Choi, H., & Claridge-Chang, A. (2019). Moving beyond P values: Data analysis with estimation graphics. Nature Methods, 16(7), 565–566. 10.1038/s41592-019-0470-3

Huffmyer, A. S., Johnson, C. J., Epps, A. M., Lemus, J. D., & Gates, R. D. (2021). Feeding and thermal conditioning enhance coral temperature tolerance in juvenile Pocillopora acuta. Royal Society Open Science, 8(5), 210644. 10.1098/rsos.210644

Hughes, T. P., Anderson, K. D., Connolly, S. R., Heron, S. F., Kerry, J. T., Lough, J. M., Baird, A. H., Baum, J. K., Berumen, M. L., Bridge, T. C., Claar, D. C., Eakin, C. M., Gilmour, J. P., Graham, N. A. J., Harrison, H., Hobbs, J. P. A., Hoey, A. S., Hoogenboom, M., Lowe, R. J., … Wilson, S. K. (2018). Spatial and temporal patterns of mass bleaching of corals in the Anthropocene. Science, 359(6371), 80–83. 10.1126/science.aan8048

Jones, R. J., & Yellowlees, D. (1997). Regulation and control of intracellular algae (= zooxanthellae) in hard corals. Philosophical Transactions of the Royal Society of London. Series B: Biological Sciences, 352(1352), 457–468. 10.1098/rstb.1997.0033

Jurriaans, S., & Hoogenboom, M. O. (2020). Seasonal acclimation of thermal performance in two species of reef-building corals. Marine Ecology Progress Series, 635, 55–70. 10.3354/meps13203

Klepac, C. N., & Barshis, D. J. (2020). Reduced thermal tolerance of massive coral species in a highly variable environment: Reduced heat tolerance of massive corals. Proceedings of the Royal Society B: Biological Sciences, 287(1933), 19–21. 10.1098/rspb.2020.1379rspb20201379

Le Nohaïc, M., Ross, C. L., Cornwall, C. E., Comeau, S., Lowe, R., McCulloch, M. T., & Schoepf, V. (2017). Marine heatwave causes unprecedented regional mass bleaching of thermally resistant corals in northwestern Australia. Scientific Reports, 7(1), Articolo 1. 10.1038/s41598-017-14794-y

Majerova, E., Carey, F. C., Drury, C., & Gates, R. D. (2021). Preconditioning improves bleaching tolerance in the reef-building coral Pocillopora acuta through modulations in the programmed cell death pathways. Molecular Ecology, 30(14), 3560–3574. 10.1111/mec.15988

Majerová, E., & Drury, C. (2022). Thermal preconditioning in a reef-building coral alleviates oxidative damage through a BI-1-mediated antioxidant response. Frontiers in Marine Science, 9. https://www.frontiersin.org/articles/10.3389/fmars.2022.971332

Marhoefer, S. R., Zenger, K. R., Strugnell, J. M., Logan, M., van Oppen, M. J. H., Kenkel, C. D., & Bay, L. K. (2021). Signatures of Adaptation and Acclimatization to Reef Flat and Slope Habitats in the Coral Pocillopora damicornis. Frontiers in Marine Science, 8. https://www.frontiersin.org/articles/10.3389/fmars.2021.704709

Martell, H. A. (2023). Thermal priming and bleaching hormesis in the staghorn coral, Acropora cervicornis (Lamarck 1816). *Journal of Experimental Marine Biology and Ecology*, *560*, 151820. 10.1016/j.jembe.2022.151820

Marzonie, M. R., Bay, L. K., Bourne, D. G., Hoey, A. S., Matthews, S., Nielsen, J. J. V., & Harrison, H. B. (2023). The effects of marine heatwaves on acute heat tolerance in corals. Global Change Biology, 29(2), 404–416. 10.1111/gcb.16473

McClanahan, T. R., Ateweberhan, M., Graham, N. a. J., Wilson, S. K., Sebastián, C. R., Guillaume, M. M. M., & Bruggemann, J. H. (2007). Western Indian Ocean coral communities: Bleaching responses and susceptibility to extinction. Marine Ecology Progress Series, 337, 1–13. 10.3354/meps337001

McClanahan, T. R., Maina, J. M., Darling, E. S., Guillaume, M. M. M., Muthiga, N. A., D’agata, S., Leblond, J., Arthur, R., Jupiter, S. D., Wilson, S. K., Mangubhai, S., Ussi, A. M., Humphries, A. T., Patankar, V., Shedrawi, G., Julius, P., Ndagala, J., & Grimsditch, G. (2020). Large geographic variability in the resistance of corals to thermal stress. Global Ecology and Biogeography, 29(12), 2229–2247. 10.1111/geb.13191

McIlroy, S. E., Cunning, R., Baker, A. C., & Coffroth, M. A. (2019). Competition and succession among coral endosymbionts. Ecology and Evolution, 9(22), 12767–12778. 10.1002/ece3.5749

Middlebrook, R., Hoegh-Guldberg, O., & Leggat, W. (2008). The effect of thermal history on the susceptibility of reef-building corals to thermal stress. Journal of Experimental Biology, 211(7), 1050–1056. 10.1242/jeb.013284

Nielsen, J. J. V., Kenkel, C. D., Bourne, D. G., Despringhere, L., Mocellin, V. J. L., & Bay, L. K. (2020). Physiological effects of heat and cold exposure in the common reef coral Acropora millepora. Coral Reefs, 39(2), 259–269. 10.1007/s00338-019-01881-x

Oliver, E. C. J., Donat, M. G., Burrows, M. T., Moore, P. J., Smale, D. A., Alexander, L. V., Benthuysen, J. A., Feng, M., Sen Gupta, A., Hobday, A. J., Holbrook, N. J., Perkins-Kirkpatrick, S. E., Scannell, H. A., Straub, S. C., & Wernberg, T. (2018). Longer and more frequent marine heatwaves over the past century. Nature Communications, 9(1), Articolo 1. 10.1038/s41467-018-03732-9

Oliver, T. A., & Palumbi, S. R. (2011). Do fluctuating temperature environments elevate coral thermal tolerance? Coral Reefs, 30(2), 429–440. 10.1007/s00338-011-0721-y

Palumbi, S. R., Barshis, D. J., Traylor-Knowles, N., & Bay, R. A. (2014). Mechanisms of reef coral resistance to future climate change. Science, 344(6186), 895–898. 10.1126/science.1251336

Putnam, H. M., & Edmunds, P. J. (2011). The physiological response of reef corals to diel fluctuations in seawater temperature. Journal of Experimental Marine Biology and Ecology, 396(2), 216–223. 10.1016/j.jembe.2010.10.026

Quigley, K. M., Randall, C. J., van Oppen, M. J. H., & Bay, L. K. (2020). Assessing the role of historical temperature regime and algal symbionts on the heat tolerance of coral juveniles. Biology Open, 9(1), bio047316. 10.1242/bio.047316

Rivest, E. B., Comeau, S., & Cornwall, C. E. (2017). The Role of Natural Variability in Shaping the Response of Coral Reef Organisms to Climate Change. Current Climate Change Reports, 3(4), 271–281. 10.1007/s40641-017-0082-x

Rodrigues, L. J., & Grottoli, A. G. (2007). Energy reserves and metabolism as indicators of coral recovery from bleaching. Limnology and Oceanography, 52(5), 1874–1882. 10.4319/lo.2007.52.5.1874

Roik, A., Wall, M., Dobelmann, M., Nietzer, S., Brefeld, D., Fiesinger, A., Reverter, M., Schupp, P. J., Jackson, M., Rutsch, M., & Strahl, J. (2023). *Trade-offs in a reef-building coral after six years of thermal acclimation* [Preprint]. Ecology. 10.1101/2023.07.20.549699

Sawall, Y., Nicosia, A. M., McLaughlin, K., & Ito, M. (2022). Physiological responses and adjustments of corals to strong seasonal temperature variations (20–28°C). Journal of Experimental Biology, 225(13), jeb244196. 10.1242/jeb.244196

Schoepf, V., Carrion, S. A., Pfeifer, S. M., Naugle, M., Dugal, L., Bruyn, J., & McCulloch, M. T. (2019). Stress-resistant corals may not acclimatize to ocean warming but maintain heat tolerance under cooler temperatures. Nature Communications, 10(1). 10.1038/s41467-019-12065-0

Schoepf, V., Jung, E. M. U., McCulloch, M. T., White, N. E., Stat, M., & Thomas, L. (2020). Differential Recovery from Mass Coral Bleaching on Naturally Extreme Reef Environments in NW Australia. MarXiv. doi:10.31230/osf.io/s9xha

Schoepf, V., Sanderson, H., & Larcombe, E. (2021). Coral heat tolerance under variable temperatures: Effects of different variability regimes and past environmental history vs. Current exposure. Limnology and Oceanography, 1–15. 10.1002/lno.12000

Schoepf, V., Sanderson, H., & Larcombe, E. (2022). Coral heat tolerance under variable temperatures: Effects of different variability regimes and past environmental history vs. Current exposure. Limnology and Oceanography, 67(2), 404–418. 10.1002/lno.12000

Schoepf, V., Stat, M., Falter, J. L., & McCulloch, M. T. (2015). Limits to the thermal tolerance of corals adapted to a highly fluctuating, naturally extreme temperature environment. Scientific Reports, 5(May), 1–14. 10.1038/srep17639

Silverstein, R. N., Cunning, R., & Baker, A. C. (2015). Change in algal symbiont communities after bleaching, not prior heat exposure, increases heat tolerance of reef corals. Global Change Biology, 21(1), 236–249. 10.1111/gcb.12706

Speelman, P. E., Parger, M., & Schoepf, V. (2023). Divergent recovery trajectories of intertidal and subtidal coral communities highlight habitat-specific recovery dynamics following bleaching in an extreme macrotidal reef environment. PeerJ, 11, e15987. 10.7717/peerj.15987

Thomas, L., Rose, N. H., Bay, R. A., López, E. H., Morikawa, M. K., Ruiz-Jones, L., & Palumbi, S. R. (2018). Mechanisms of Thermal Tolerance in Reef-Building Corals across a Fine-Grained Environmental Mosaic: Lessons from Ofu, American Samoa. Frontiers in Marine Science, 4. https://www.frontiersin.org/articles/10.3389/fmars.2017.00434

Van Oppen, M. J. H., Oliver, J. K., Putnam, H. M., & Gates, R. D. (2015). Building coral reef resilience through assisted evolution. Proceedings of the National Academy of Sciences of the United States of America, 112(8), 2307–2313. 10.1073/pnas.1422301112

Voolstra, C. R., Buitrago-López, C., Perna, G., Cárdenas, A., Hume, B. C. C., Rädecker, N., & Barshis, D. J. (2020). Standardized short-term acute heat stress assays resolve historical differences in coral thermotolerance across microhabitat reef sites. Global Change Biology, 26(8), 4328–4343. 10.1111/gcb.15148

Voolstra, C. R., & Ziegler, M. (2020). Adapting with Microbial Help: Microbiome Flexibility Facilitates Rapid Responses to Environmental Change. BioEssays, 42(7), 2000004. 10.1002/bies.202000004

Wall, M., Doering, T., Pohl, N., Putchim, L., Ratanawongwan, T., & Roik, A. (2023). Natural thermal stress-hardening of corals through cold temperature pulses in the Thai Andaman Sea (p. 2023.06.12.544549). bioRxiv. 10.1101/2023.06.12.544549

Wiedenmann, J., D’Angelo, C., Mardones, M. L., Moore, S., Benkwitt, C. E., Graham, N. A. J., Hambach, B., Wilson, P. A., Vanstone, J., Eyal, G., Ben-Zvi, O., Loya, Y., & Genin, A. (2023). Reef-building corals farm and feed on their photosynthetic symbionts. Nature, 620(7976), Articolo 7976. 10.1038/s41586-023-06442-5

Williamson, O. M., Allen, C. E., Williams, D. E., Johnson, M. W., Miller, M. W., & Baker, A. C. (2021). Neighboring colonies influence uptake of thermotolerant endosymbionts in threatened Caribbean coral recruits. Coral Reefs, 40(3), 867–879. 10.1007/s00338-021-02090-1

Ziegler, M., Seneca, F. O., Yum, L. K., Palumbi, S. R., & Voolstra, C. R. (2017). Bacterial community dynamics are linked to patterns of coral heat tolerance. Nature Communications, 8, 1–8. 10.1038/ncomms14213

